# Targeted tumor desialylation by engineered probiotic reprograms the tumor microenvironment to potentiate immunotherapy

**DOI:** 10.64898/2026.10.04.756258

**Authors:** Jinhui Wang, Jing Wang, Tao OuYang, Bin Shi, He Meng, Zong-Wan Mao, Wei Xia

## Abstract

Tumor hypersialylation drives immunosuppression and limits immunotherapy efficacy by shielding tumor cells and engaging inhibitory immune receptors. Here, we engineered *Escherichia coli* Nissle 1917 (EcN^Δpgi^-SAS) as a metabolically confined “desialylation catalyst” to remodel the tumor microenvironment. This strain combines HER2-directed adhesion, hypoxia-inducible surface sialidase display for targeted glycan cleavage, and sialic acid catabolism to prevent glycolytic recycling. Critically, disrupting glucose metabolism (Δ*pgi*) restricts bacterial proliferation to sialic acid-rich tumor microenvironment, significantly enhancing biosafety. In mice, EcN^Δpgi^-SAS colonized tumors and effectively desialylated tumor cells, synergizing with anti-PD-L1 to suppress growth of immunologically “cold” tumors. Mechanistically, desialylation reversed immunosuppression by broadly enhancing antitumor immunity. Orally administered EcN^Δpgi^-SAS substantially inhibited colorectal carcinogenesis, repaired epithelial barrier integrity by restoring tight junctions, and prevented adenoma formation. This integrated strategy establishes a clinically translatable platform for reversing immunosuppression in solid tumors.

## Introduction

Cancer immunotherapy has revolutionized clinical tumor treatment. Particularly, immune checkpoint blockade targeting PD-1, PD-L1, and CTLA-4 has successfully eradicated metastatic tumors in certain patients by reactivating endogenous anti-tumor immunity ^1-4^. However, the majority of patients exhibit no responses, relapse or even resistance to treatment, which is primarily attributed to the immunosuppressive tumor microenvironment (TME). The TME constitutes a major therapeutic obstacle that compromises immune cell infiltration, inhibits anti-tumor immune activation, and drives broad immunotherapy resistance in most solid tumors ^5-7^. Recent evidence implicates cell-surface glycosylation as an important mediator of immune suppression ^8-10^. Notably, tumor hypersialylation emerged as a prevalent and conserved mechanism driving TME immunosuppression ^11-14^. Pathologically elevated sialylation forms a dense, negatively charged glycocalyx barrier on tumor cell surfaces that masks antigen epitopes, generates spatial steric hindrance, and impedes antigen recognition, dendritic cell (DC) cross-presentation and T-cell immunological synapse formation ^15-17^. Furthermore, abundant sialic acid residues on tumor cell and immune cell surfaces engage inhibitory Siglec receptors, suppressing cytotoxic lymphocyte function, polarizing macrophages toward immunosuppressive phenotypes, and establishing a persistent immune-evasive niche ^18-21^. To overcome sialylation-mediated immunosuppression, targeted desialylation strategies employing sialidase-conjugated therapeutics, including antibody-sialidase conjugates and bispecific T-cell engager (BiTE)-sialidase fusion proteins, have been developed to selectively remove tumor surface sialoglycans and restore anti-tumor immunity ^22-25^. Despite promising preclinical activity, these synthetic biologics suffer from inherent limitations, including systemic instability, off target desialylation, and limited therapeutic durability ^22, 26^. Most critically, sialic acid liberated by exogenous sialidases can be recycled and reused by tumor cells to rapidly restore surface hypersialylation, forming a compensatory glycan remodeling loop that substantially compromises long-term immunotherapeutic efficacy ^27, 28^.

Engineered bacteria have emerged as potent living therapeutics for tumor intervention due to their intrinsic tumor tropism, in situ payload delivery capability, and programmable genetic versatility ^29-31^. As a well-characterized probiotic strain, *Escherichia coli* Nissle 1917 (EcN) colonizes hypoxic tumor tissues and demonstrates favorable biocompatibility in vivo. To date, most EcN-based anti-tumor strategies primarily utilize EcN as a passive delivery vehicle to express and release immunomodulatory effectors, such as nanobodies, cytokines and neoantigen, which amplify systemic anti-tumor immune responses ^32-38^. In contrast, much fewer studies repurpose EcN as an active catalytic chassis to directly remodel the immunosuppressive TME ^39-42^. Despite these advantages, conventional engineered EcN strains exhibit unconstrained vivo proliferation, potentially causing systemic bacterial dissemination, local inflammation, and off-target toxicity. This inherent biosafety risk remains a major translational barrier for clinical applicability of EcN-based tumor therapies ^43, 44^.

Here, we constructed a multifunctional, biosafely optimized EcN strain, EcN^Δpgi^-SAS. This engineered bacterium integrates tumor targeting, hypoxia-inducible catalytic desialylation, sialic acid metabolic scavenging, and tumor-restricted proliferation. By combining HER2 affibody-mediated tumor anchoring and hypoxia-controlled sialidase expression, the strain achieves highly specific and localized tumor desialylation. Concurrently, an embedded sialic acid catabolic cluster irreversibly eliminates liberated sialic acid to block tumor glycan recycling. Furthermore, disruption of the phosphoglucoisomerase gene (*pgi*) restricts bacteria to sialic acid as the sole carbon source, thereby limiting bacterial proliferation exclusively within sialic acid-rich tumor tissues. This genomic modification mitigates systemic bacterial toxicity and substantially improves in vivo biosafety. We demonstrated that EcN^Δpgi^-SAS remodels the immunosuppressive TME, converting immunologically “cold” tumors into “hot” tumors, and synergizes with PD-L1 antibody to suppress subcutaneous tumor growth in mice. Notably, oral administered EcN^Δpgi^-SAS achieves substantial therapeutic efficacy in AOM/DSS-induced colorectal cancer models and repairs damaged intestinal epithelial barriers. This study establishes a novel targeted-desialylation live microbial immunotherapy strategy, offering a safe, durable, and clinically translatable approach for solid tumor intervention.

## Results

### Design and construction of engineered bacteria with desialylating and metabolic activities

To enable the probiotic EcN to deplete tumor sialoglycans, we aimed to engineer EcN to express a sialidase enzyme on its surface. *Streptococcus pneumoniae* (*S. pneumonia*) sialidase A (*Sp*NanA) is one of the most extensively studied bacterial sialidase, known for its potent desialylation activity ^39^. We first assessed recombinant *Sp*NanA activity in vitro using glycoprotein fetuin as substrate. Incubation *Sp*NanA with fetuin at a 1:100 ratios at 37°C for 1 h resulted in efficient removal of sialic acids on fetuin, evidenced by a distinct molecular weight shift on SDS-PAGE **(Supplementary Fig. 1a)**. This shift was absent when using the catalytically inactive mutant *Sp*NanA^E647A^ mutant **(Supplementary Fig. 1b)**. MALDI-TOF mass spectrometry confirmed sialic acid removal of *Sp*NanA-treated fetuin **(Supplementary Fig. 2)**. To evaluate the desialylation activity of *Sp*NanA toward cells, we treated a panel of cancer cell lines (HeLa, MCF7, SK-BR-3, A549, MC38, AGS, and 4T1) with purified *Sp*NanA at 37°C for 30 min. Subsequent *Sambucus nigra* agglutinin (SNA-Cy5) staining revealed significant reduction of surface sialoglycans across all tested lines **(Supplementary Fig. 3)**. Collectively, these results demonstrated robust desialylation activity of *Sp*NanA against both glycoproteins and tumor cells.

Subsequently, we engineered EcN for surface display of *Sp*NanA to facilitate direct access of the sialidase to tumor cell membrane sialoglycans, thereby maximizing catalytic efficiency through proximity effects. We constructed a stable plasmid pSYSU-1 encoding an Lpp-OmpA-based surface display system for *Sp*NanA ^45^. In this construct, *Sp*NanA (54 kDa) was fused C-terminally to the bacterial lipoprotein OmpA (16 kDa) under control of the hypoxia-responsive P*fnrS* promoter to enable anaerobically inducible expression. To enhance tumor targeting, an anti-HER2 affibody (ZHER2)-AIDA surface display cassette, governed by the constitutive P*lacUV5* promoter, was incorporated into the same plasmid to confer HER2-binding capacity. Previous studies demonstrate that tumor cells scavenge free sialic acid from their microenvironment for reuse, maintaining high surface sialylation levels essential for immune evasion and metastasis ^28, 46^. To prevent tumor recycling of liberated sialic acid by sialidase, we constructed another stable plasmid pSYSU-2 containing a *E. coli* sialic acid catabolism gene cluster (*nanT, nanA, nanK, nanE, nagA, nagB*) ^47^ under the control of the sialic acid-responsive P*nanR* promoter. This system facilitates bacterial to import of sialic acid via the NanT transporter and sequential degradation of sialic acid into fructose-6-phosphate through concerted enzymatic catalysis, which feeds irreversibly into bacterial glycolysis and ultimately the tricarboxylic acid (TCA) cycle. (**Fig. 1a and Supplementary Fig. 4)**. The probiotic strain EcN-SAS was engineered by co-transforming probiotic EcN with the two stable plasmids: one encoding the *Sp*NanA sialidase (S) and anti-HER2 affibody surface display system (A), and the other containing the sialic acid catabolism gene clusters (S).

**Figure 1.**
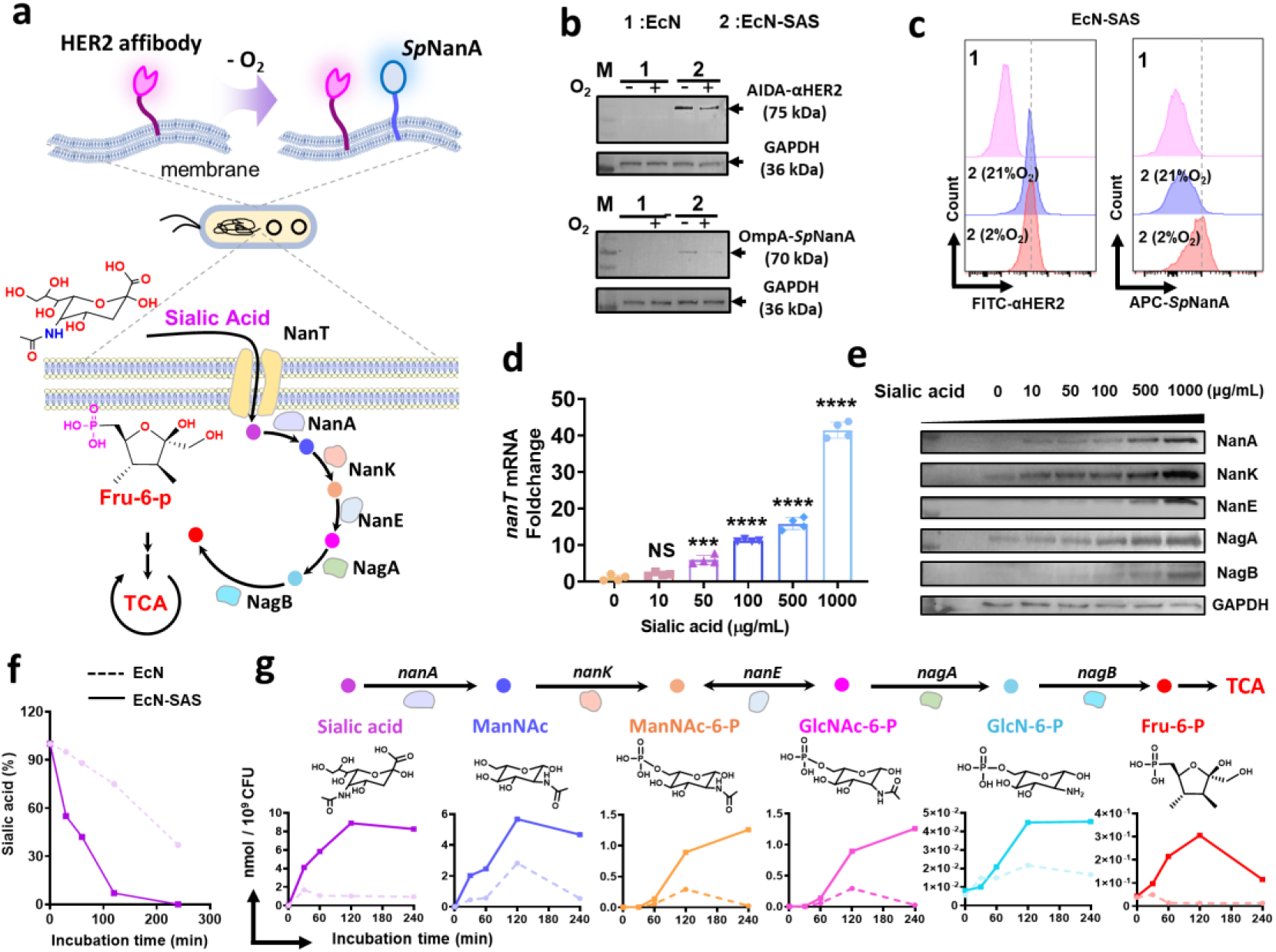
Construction and characterization of engineered bacteria for tumor-targeted desialylation. **a**, Schematic illustration of engineered *Escherichia coli* Nissle 1917 (EcN) strains harboring two stable plasmids pSYSU-1 and pSYSU-2. pSYSU-1 contains a hypoxia-inducible P*fnrS* promoter driving expression of *ompA-nanA* fusion gene (encoding OmpA-*Sp*NanA) and a constitutive P*lacuv5* promoter driving the gene encoding AIDA-anti-HER2 affibody (αHER2). pSYSU-2 contains a constitutively expressed *nanR* gene and a sialic acid-inducible P*nanR* promoter governing the *nanT-nanA-nanK-nanE-nagA-nagB* catabolic cluster. **b**, Western blot detection of AIDA-αHER2 (75 kDa) and OmpA-*Sp*NanA (70 kDa) expression in wild-type EcN and engineered EcN-SAS under normoxia (21% O2) and hypoxia (2% O2). GAPDH served as a loading control. **c**, Flow cytometry quantification of surface-displayed αHER2 and *Sp*NanA under normoxia and hypoxia. **d**, qPCR analysis of *nanT* transcriptional induction in response to exogenous sialic acid. Data are presented as mean ± s.d. from n = 4 biological replicates. Statistical analysis was performed using one-way ANOVA followed by Dunnett’s multiple comparisons test \*\*\**P* < 0.001 and \*\*\*\**P* < 0.0001. NS, not significant. **e**, Western blot detection of NanA, NanK, NanT, NagA and NagB expression after incubation with different sialic acid concentrations. **f**, Kinetics of extracellular sialic acid catabolism. EcN-SAS or a EcN control were cultured in medium supplemented with 2 mM sialic acid, and residual sialic acid in the supernatants were quantified by LC-MS at indicated time. **g**, Intracellular levels of sialic acid and catabolic intermediates in EcN-SAS relative to EcN control.

Western blot analysis revealed constitutive expression of the ZHER2-AIDA fusion protein under both normoxic and hypoxic conditions. In contrast, the OmpA-*sp*NanA fusion protein was detectable only under hypoxic conditions (**Fig. 1b**). Flow cytometry confirmed these findings, demonstrating that ZHER2 was displayed on the surface of engineered EcN-SAS bacteria under both oxygen states, while *Sp*NanA surface display occurred exclusively under hypoxia (**Fig. 1c**). The sialidase activity of surface-displayed *Sp*NanA on engineered bacteria was subsequently assessed using a fluorogenic assay kit. In brief, varying amounts of EcN-SAS were incubated with a fluorogenic sialic acid-conjugated substrate under hypoxic conditions. Time-course fluorescence measurements demonstrated that the substrate cleavage rate increased proportionally with bacterial density, confirming the potent activity of the surface-displayed sialidase (**Supplementary Fig. 5**). To evaluate the HER2 receptor-binding capability of the surface-displayed ZHER2 affibody, EcN-SAS with a genomically integrated superfold GFP (sfGFP) reporter was incubated with murine 4T1 cells or human HER2-transfected 4T1 cells (4T1-HER2) for 4 h. After extensive washing, substantially higher adherence of EcN-SAS was observed on 4T1-HER2 compared to 4T1 cells, confirming successful functionalization of the ZHER2 affibody (**Supplementary Fig. 6**).

Expression analysis of the sialic acid catabolism gene cluster demonstrated concentration-dependent induction of cluster transcription and NanA, NanK, NanT, NagA, and NagB protein expression by exogenous sialic acid, as evidenced by quantitative PCR (qPCR) and western blot (**Fig. 1d, e**). To assess metabolic functionality, EcN-SAS was cultured in medium containing 2 mM sialic acid. Liquid chromatography-mass spectrometry (LC-MS) quantification of culture supernatants revealed near-complete substrate consumption within 120 minutes, whereas a sialic acid catabolism-deficient strain (EcN-SA) metabolized approximately 30% during this period (**Fig. 1f**). Intracellular metabolite profiling showed significantly elevated levels of sialic acid and metabolism intermediates in EcN-SAS compared to EcN-SA (**Fig. 1g**), confirming efficient sialic acid import and metabolic flux toward fructose-6-phosphate conversion. Collectively, these results establish that EcN-SAS integrates HER2 receptor targeting, hypoxia-inducible sialidase display, and a high-efficiency sialic acid catabolic system.

### Engineered bacteria selectively desialylates HER2^+^ tumor cells and catabolizes liberated sialic acids

We subsequently assessed the targeted desialylation capability and sialic acid catabolism properties of the engineered bacteria at the cellular level. EcN-SAS was co-incubated with 4T1 or 4T1-HER2 tumor cells at varying bacterium-to-tumor cell ratios under hypoxic conditions for 3 h. Post-incubation, cell surface sialylation was evaluated via flow cytometry. At a low ratio (10:1, bacteria: cells), sialic acid removal was markedly higher on 4T1-HER2 cells compared to 4T1 cells, confirming enhanced targeting specificity conferred by HER2 binding (**Fig. 2a, 2b)**. Time-course analysis revealed that EcN-SAS reduced sialic acids on 4T1-HER2 cells (10:1, EcN-SAS: 4T1-HER2 cells) non-detectable levels within 200 min, whereas a sialidase catalytically inactive strain EcN-S^E647A^AS exhibited no desialylation capability (**Fig. 2c**). To further visualize targeted tumor cell desialylation, both EcN-SAS and an isogenic ZHER2-deficient bacterial strain (EcN-SS) were genomically integrated with a sfGFP reporter for fluorescence detection. These engineered bacteria were co-cultured with four tumor cell lines, murine 4T1, 4T1-HER2, human HER2^+^ SK-BR-3 and HER2^-^ MDA-MB-231 under normoxic or hypoxic conditions. After incubation, tumor cells were washed and bacterial-tumor cell interactions were analyzed by confocal microscopy (**Fig. 2d**). In consistent, substantially higher numbers of EcN-SAS compared to EcN-SS were observed tightly adhering to HER2^+^ cells (4T1-HER2 and SK-BR-3). Critically, EcN-SAS-mediated desialylation was strictly hypoxia-dependent, as enzymatic activity occurred exclusively under hypoxic conditions (**Fig. 2e-f**). In contrast, HER2^-^ cells (4T1 and MDA-MB-231) exhibited minimal bacterial adhesion and no detectable desialylation, further confirming HER2-dependent targeting specificity **(Supplementary Fig. 7-8)**.

**Figure 2.**
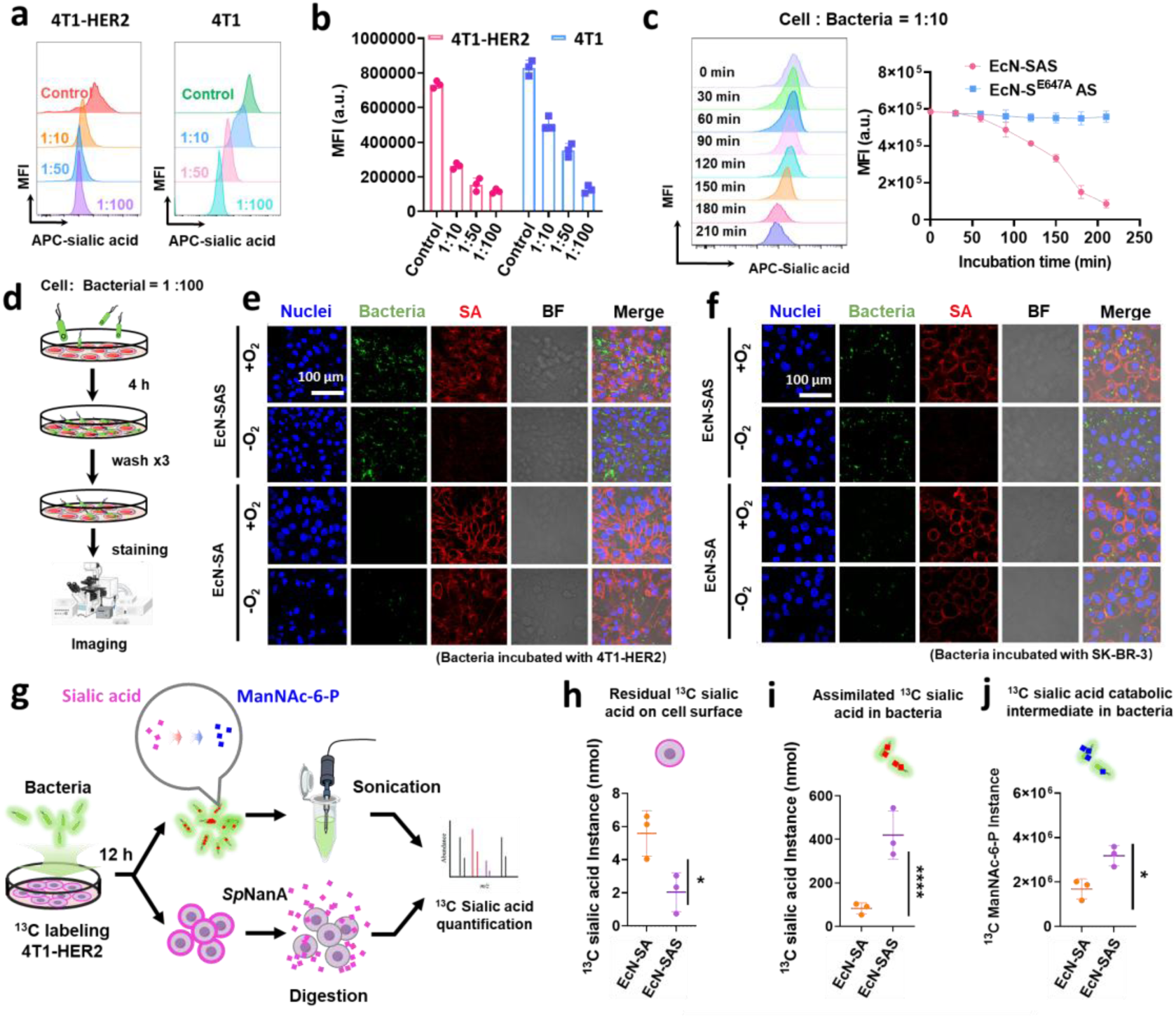
Engineered bacteria mediate targeted desialylation of HER2^+^ tumors and metabolic consumption of liberated sialic acids. **a,** Flow cytometry profiles of human HER2-transfected 4T1 (4T1-HER2) and 4T1 cells stained with Sambucus nigra agglutinin (SNA-Cy5) after co-culture with engineered EcN-SAS strain at indicated tumor cell-to-bacteria ratios. **b**, Quantification of cell-surface sialylation (SNA-Cy5) on HER2^+^ 4T1-HER2 and HER2^-^ 4T1 cells post co-culture with EcN-SAS. **c**, Desialylation kinetic of 4T1-HER2 cells by EcN-SAS over 240 min. **b, c**, Data are presented as mean ± s.d. from n = 3 biological replicates. **d**, Schematic workflow for confocal microcopy assessment of bacteria-mediated tumor desialylation. **e, f**, Confocal images of bacteria-mediated tumor-targeting desialylation. Engineered EcN-SAS or EcN-SA was co-cultured with 4T1-HER2 (**e**) or HER2^+^ SK-BR-3 cells (**f**) under normoxia (21% O2) and hypoxia (2% O2) for 4 h. Blue: nuclei; red: sialic acids; green: engineered bacteria. Scale bar, 100 μm. **g**, Schematic workflow of ^13^C-labeled sialic acid competition assay. ^13^C-sialic acid labeled 4T1-HER2 was co-incubated with engineered EcN-SAS or EcN-SA strains. Quantification of tumor cell-surface residual ^13^C-sialylation levels (**h**), intracellular bacterial ^13^C-sialic acid (**i**) and bacterial catabolic intermediate ^13^C-ManNAc-6-P (**j**). Data are presented as mean ± s.d from n = 3 biological replicates. Statistical analysis was performed using unpaired Student’s t-test. \**P* < 0.05 and \*\*\*\**P* < 0.0001.

Previous studies have demonstrated that tumor cells utilize free sialic acids in TME for surface sialylation. To determine whether the engineered bacteria could compete with tumor cells for free sialic acids and subsequently degrade them, we metabolically labelled 4T1-HER2 cells with ^13^C-sialic acid. These labeled tumor cells were co-culture with either EcN-SAS or EcN-SA (lacking sialic acid catabolism gene cluster) under hypoxic conditions for 12 h. The engineered bacteria were washed three times with PBS, collected, and lysed via sonication. Subsequently, the 4T1-HER2 tumor cells were further washed extensively with PBS to eliminate adherent bacteria. Surface sialylation of tumor cells was stripped using excess sialidase to liberate free sialic acids. Free ^13^C-sialic acid from bacterial lysates and stripped tumor cells was subsequently quantified by LC-MS in selected ion monitoring (SIM) mode (**Fig. 2g**). Tumor cells co-cultured with EcN-SAS retained significantly less surface ¹³C-sialic acid than those exposed to EcN-SA, which lacks functional sialic acid catabolism gene clusters (**Fig. 2h**). Concurrently, bacterial lysates of EcN-SAS exhibited markedly higher ¹³C-sialic acid levels than those of EcN-SA (**Fig. 2i**). Furthermore, EcN-SAS lysates contained substantially elevated levels of ¹³C-labeled ManNAc-6P, a specific metabolite generated by the NanA-NanK enzymatic cascade (**Fig. 2j**). Collectively, these results indicate that engineered bacteria equipped with sialic acid catabolism gene clusters compete with tumor cells to internalize free sialic acid, whereupon the scavenged sialic acid enters the bacterial catabolic pathway.

### Generation of an attenuated EcN strain for tumor-specific proliferation and improved in vivo biosafety

The major risk of bacteria-based therapy is infection-associated toxicity and systemic pathogenesis, wherein uncontrolled bacterial proliferation may cause severe complications like sepsis, healthy tissue damage or multi-organ failure ^48, 49^. To improve biosafety during tumor therapy, we sought to engineer an attenuated EcN-SAS strain through targeted metabolic disruption. EcN-SAS can internalize free sialic acids and degrade them through the NanA-NanK enzymatic cascade, which ultimately generates fructose-6-phosphate to fuel the bacterial TCA cycle and support proliferation. Based on this metabolic feature, we hypothesized that disruption of glycolysis via knockout of the phosphoglucose isomerase gene (*pgi*) would render bacterial growth strictly dependent on sialic acid catabolism. Since phosphoglucose isomerase catalyzes the conversion of glucose-6-phosphate to fructose-6-phosphate, this genetic design forces bacteria to rely exclusively on sialic acid-derived fructose-6-phosphate as a carbon source. Accordingly, the engineered strain proliferates robustly in sialic acid-rich TME but exhibits limited growth in sialic acid-deficient normal tissues (**Fig. 3a**). Using a previously reported CRISPR/Cas9 strategy ^50^, we knocked out the *pgi* gene in the EcN-SAS to generate the attenuated strain EcN^Δ^*^pgi^-*SAS. When cultured in LB medium, EcN^Δ^*^pgi^*-SAS exhibited markedly slower growth than EcN-SAS. Glucose supplementation substantially promoted the growth of EcN-SAS, but had no notable effect on EcN^Δ^*^pgi^*-SAS. In contrast, supplementation with 5 mM fructose-6-phosphate fully restored the growth of EcN^Δ^*^pgi^*-SAS to levels comparable to EcN-SAS. These results confirmed successful blockade of the glucose-6-phosphate to fructose-6-phoshpate conversion in EcN^Δ^*^pgi^*-SAS (**Fig. 3b and Supplementary Fig. 9)**. Notably, the growth deficiency of EcN^Δ^*^pgi^*-SAS strain could also be rescued by addition of 5 mM sialic acid. The distinct growth discrepancies of EcN^Δ^*^pgi^*-SAS between sialic acid-rich and sialic acid-poor conditions demonstrated that this attenuated strain can recover from growth restriction specifically within the TME (**Fig. 3c**).

**Figure 3.**
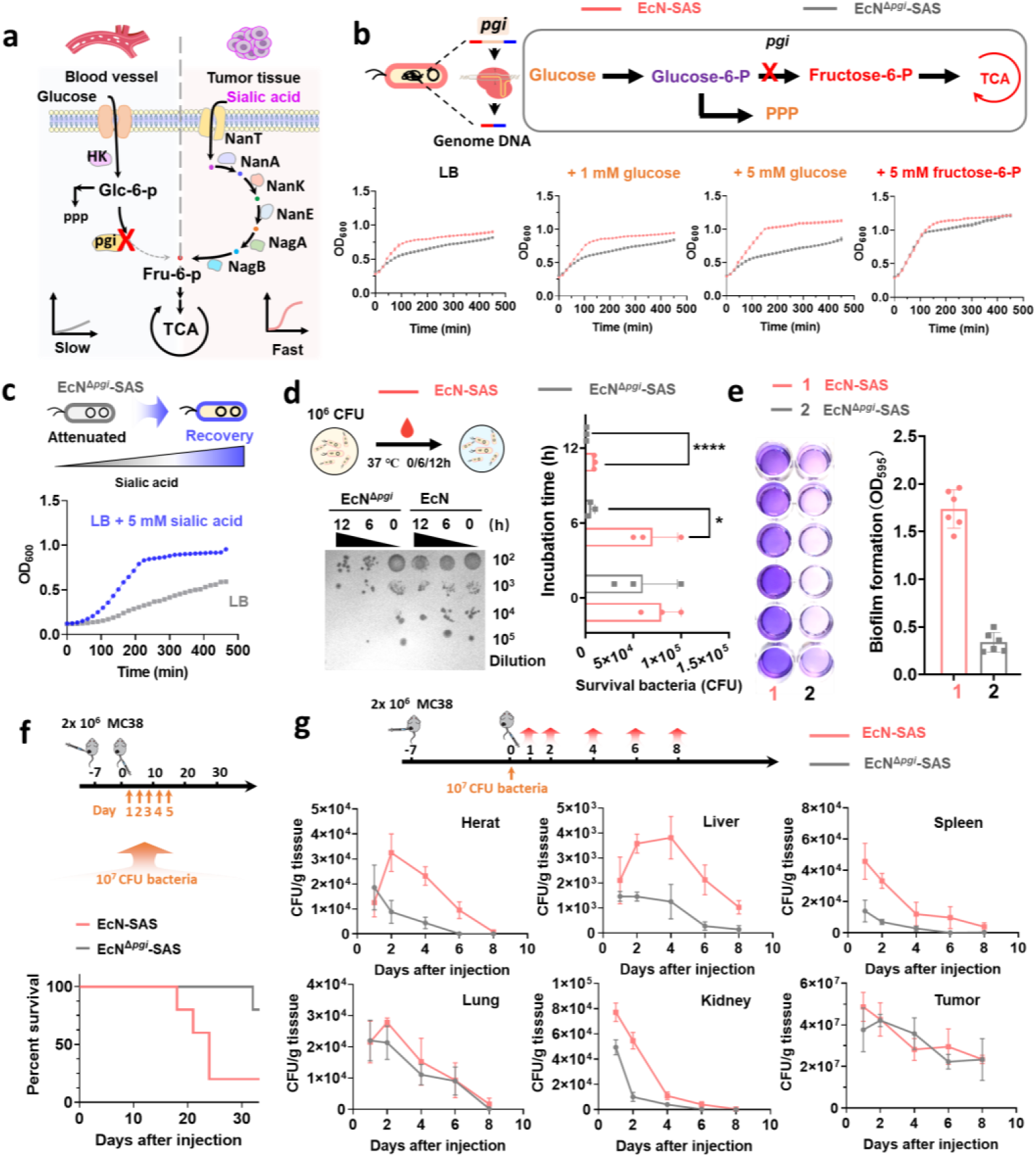
Construction of attenuated bacteria for tumor-specific proliferation and improved in vivo biosafety. **a**, Schematic illustration of bacterial attenuation strategy. Bacterial glycolysis was disrupted by deletion the phosphoglucose isomerase gene (*pgi*). The bacteria exhibited attenuated growth during systemic circulation, yet regain proliferative capacity within tumor tissue by utilizing sialic acid as a carbon source. **b**, Growth curves of EcN-SAS and attenuated EcN^Δ*pgi*^-SAS cultured in LB medium supplemented with 1 mM glucose, 5 mM glucose or 5 mM fructose-6-phosphate. EcN^Δ*pgi*^-SAS exhibited impaired growth in LB medium, growth was restored by fructose-6-phosphate but not glucose. **c**, Growth rescue of EcN^Δ*pgi*^-SAS upon supplementation with 5 mM sialic acid. **d**, Comparison of the sensitivity of EcN-SAS and EcN^Δ*pgi*^-SAS to blood-mediated clearance. Data are presented as mean ± s.d. from n = 3 biological replicates. Statistical analysis was performed using unpaired Student’s t-test. \**P* < 0.05 and \*\*\*\**P* < 0.0001. **e**, Comparison of biofilm formation capability between EcN-SAS and EcN^Δ*pgi*^-SAS. Data are presented as mean ± s.d. from n = 6 biological replicates. **f**, Survival curves of MC38 tumor-bearing mice receiving intravenous administration of 1 × 10⁷ CFU EcN-SAS or EcN^Δ*pgi*^-SAS every two days for five total doses. **g**, Bacterial colonization in heart, liver, spleen, kidney and tumor at the indicated time points following a single intravenous injection of 1 × 10⁷ CFU EcN-SAS or EcN^Δ*pgi*^-SAS. Data are presented as mean ± s.d. from n = 5 biologically independent mice per group.

As in vivo persistence and adhesion capacity constitute critical biosafety indicators for genetically engineered bacteria, we evaluated the susceptibility of EcN^Δ^*^pgi^*-SAS to blood clearance and its biofilm-formation capability. Incubation in human blood over varying durations revealed that EcN^Δ^*^pgi^*-SAS exhibited approximately 100-fold higher sensitivity to blood clearance relative to WT EcN (**Fig. 3d**). Moreover, EcN^Δ^*^pgi^*-SAS also exhibited significantly attenuated capability in biofilm formation, a key phenotypic property that enables bacteria to resist host immune clearance and antimicrobial stress (**Fig. 3e**) ^51^. Subsequently, we assessed the in vivo biosafety of EcN^Δ^*^pgi^*-SAS using tumor-bearing mouse models. In brief, MC38 tumor-bearing C57BL/6 mice were intravenously injected with 10⁷ CFU of either EcN^Δ*pgi*^-SAS or its parental strain EcN-SAS once every two days, for a total of five administrations. Mice treated with the attenuated EcN^Δ^*^pgi^*-SAS achieved an 80% survival rate after 30 days, which was markedly superior to the 20% survival rate observed in mice receiving the EcN-SAS treatment (**Fig. 3f**). To further assess the in vivo biosafety of EcN^Δ^*^pgi^*-SAS, we harvested tissues from multiple organs in mice at 2-day intervals following a single intravenous administration of 10⁷ CFU of either EcN^Δ^*^pgi^*-SAS or EcN-SAS strain. Bacterial quantification demonstrated substantially reduced colonization in the heart, liver, spleen, and kidneys at all sampled time points in mice administered EcN^Δ^*^pgi^*-SAS compared to those receiving EcN-SAS treatment. In contrast, both strains exhibited high and comparable levels of colonization within tumors (**Fig. 3g and Supplementary Fig. 10)**. Collectively, all these results demonstrated that *pgi* gene knockout substantially attenuates the systemic toxicity of the engineered bacteria, improves their biosafety for in vivo administration, yet preserves their ability to selectively accumulate in tumor tissues.

### Tumor cell desialylation by engineered bacteria potentiates immune cell-mediated clearance

As hypersialylation on tumor cells have been reported to attenuate immune cell activation and effector function by recruiting sialic acid-binding immunoglobulin-like lectins (Siglecs) ^52, 53^. We subsequently investigated whether desialylation of tumor cells by engineered bacteria EcN^Δ^*^pgi^*-SAS may sensitize them to immune cells-mediated killing. To evaluate the impact of specific bacterial functional modules on tumor cell desialylation, additional two isogenic strains of EcN^Δ^*^pgi^*-SAS were constructed, including EcN^Δ^*^pgi^*-SS (attenuated strain displaying active sialidase and with sialic acid catabolism capability) and EcN^Δ^*^pgi^*-S^E647A^S (attenuated strain displaying inactive sialidase and with sialic acid catabolism capability). In brief, HER2^+^ SK-BR-3 or HER2^-^ MDA-MB-231 cells were pre-incubated with engineered bacteria at a 1:10 ratios for 4 hours. Following extensive PBS washing to remove bacteria, fresh isolated human peripheral blood mononuclear cells (PBMCs) or NK-92 cells were added to the pre-treated SK-BR-3 or MDA-MB-231 cells at a 5:1 ratio and co-cultured for 4 hours. Immune cell-mediated tumor killing was quantified by lactate dehydrogenase (LDH) release assay. Compared to the sialidase-inactive strain EcN^Δ^*^pgi^*-S^E647A^S, EcN^Δ^*^pgi^*-SS significantly augmented PBMC-mediated killing of HER2^+^ SK-BR-3 cells, and this effect was further amplified by the HER2-targeting strain EcN^Δ^*^pgi^*-SAS (**Fig. 4a**). In contrast, HER2-targeting conferred no detectable benefit for HER2^-^ MDA-MB-231 cells. Parallel results were observed with NK-92 cells (**Fig. 4b**).

**Figure 4.**
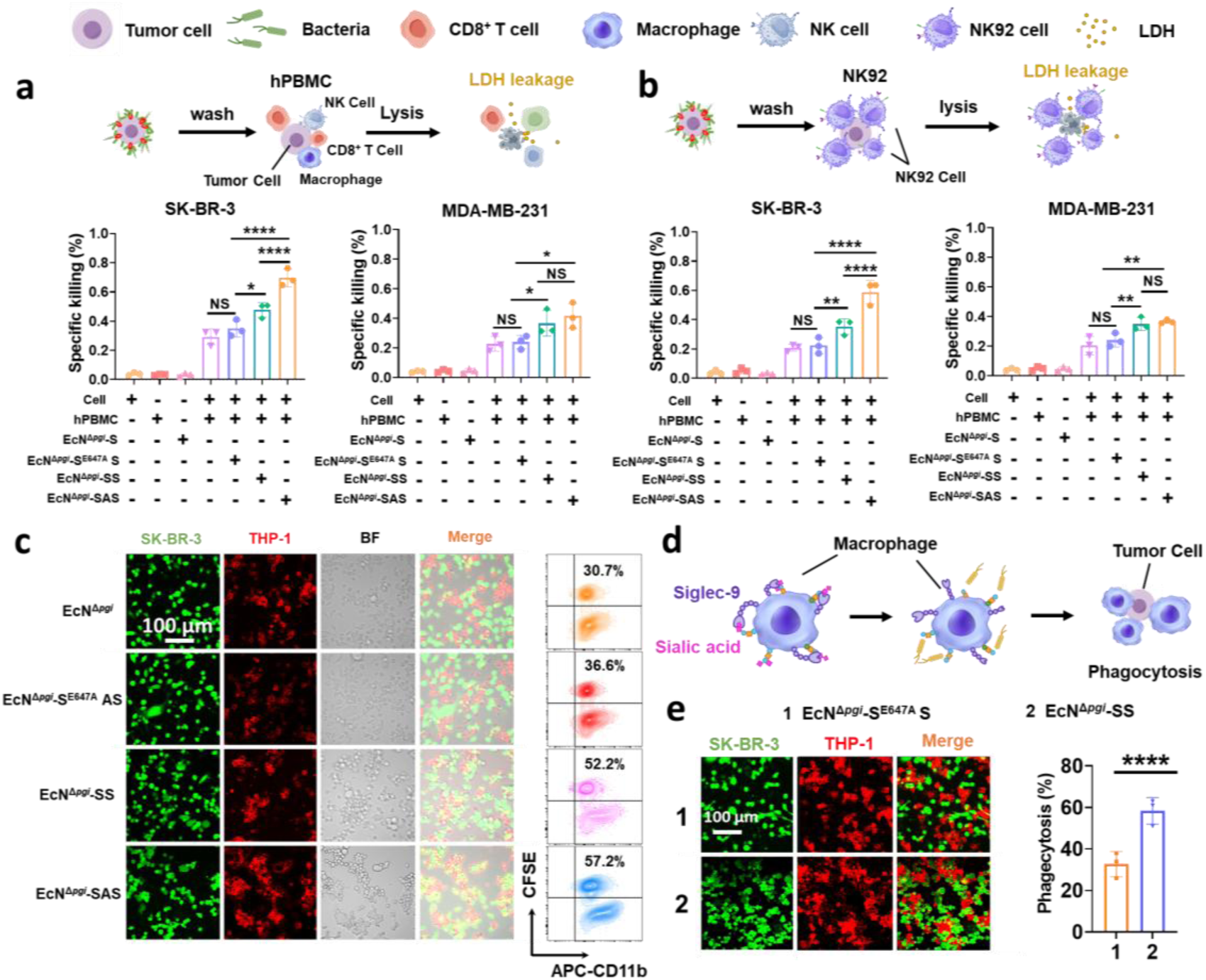
Engineered bacteria enhance immune-mediated tumor eradication via targeted desialylation. **a, b**, human peripheral blood mononuclear cells (hPBMC) (**a**) or NK92 (**b**)-mediated cytotoxicity against HER2⁺ SK-BR-3 or HER2⁻ MDA-MB-231 tumor cells pre-treated with the indicated engineered bacterial strains. Data are presented as mean ± s.d. from n = 3 biological replicates. Statistical analysis was performed using one-way ANOVA followed by Tukey’s multiple comparisons test. \**P* < 0.05, \*\**P* < 0.01 and \*\*\*\**P* < 0.0001. NS, not significant. **c**, Confocal images of THP-1-derived macrophage phagocytosis of SK-BR-3 tumor cells pre-treated with the indicated engineered bacteria strain. Phagocytosis rates were quantified as the percentage of CFSE⁺ macrophages that were CD11b⁺. **d**, Schematic workflow for macrophage phagocytosis assay after bacterial desialylation. **e**, Confocal images and quantitative analysis of tumor phagocytosis by THP-1-derived macrophage pre-treated with EcN^Δ*pgi*^-S^E647A^ or EcN^Δ*pgi*^-S strains. Data are presented as mean ± s.d. from n = 3 biological replicates. Statistical analysis was performed using unpaired Student’s t-test. \*\*\*\**P* < 0.0001.

Previous studies have demonstrated that tumor-derived sialylation drives the polarization of tumor-associated macrophages (TAMs) toward a protumorigenic M2-like phenotype via Siglec receptor binding ^25, 54^. We therefore examined the macrophage-mediated phagocytosis of tumor cells after treatment with various engineered bacteria. HER2^+^ SK-BR-3 cells were treated with EcN^Δ^*^pgi^*, EcN^Δ^*^pgi^*-S^E647A^S, EcN^Δ^*^pgi^*-SS or EcN^Δ^*^pgi^*-SAS as described above. Human THP-1-derived macrophages were then added to the pre-treated SK-BR-3 cells at a 5:1 ratio and co-cultured for 8 hours. Carboxyfluorescein succinimidyl ester (CFSE)-labeled tumor cells and macrophages stained with APC-conjugated anti-mouse CD11b antibody were subject to phagocytosis analysis. Immunofluorescence imaging revealed substantially enhanced phagocytosis of tumor cells following treatment with EcN^Δ^*^pgi^*-SS or EcN^Δ^*^pgi^*-SAS relative to treatment with EcN^Δ^*^pgi^* or the sialidase-inactive EcN^Δ^*^pgi^*-S^E647A^S strains. Flow cytometry was further employed to quantify tumor cells phagocytosis rates. Consistent with imaging results, flow cytometry analysis confirmed that tumor cell incubated with EcN^Δ*pgi*^-SAS yielded the highest phagocytosis rate of 57.2%, while those treated with EcN^Δ^*^pgi^*-SS showed a moderately reduced phagocytosis rate of 52.2%. In contrast, tumor cells exposed to EcN^Δ^*^pgi^* or EcN^Δ^*^pgi^*-S^E647A^S exhibited substantially lower phagocytosis rates of 30.7% and 36.6%, respectively (**Fig. 4c**). Similar results were observed with primary mouse bone marrow-derived macrophages (BMDMs) and RAW 264.7 cells **(Supplementary Fig. 11-12**). Collectively, these findings demonstrate that engineered bacteria-mediated tumor desialylation potentiates immune-mediated tumor elimination, while HER2-targeting specifically amplifies tumor desialylation.

Beyond modulation by tumor-derived sialic acids, macrophage-surface sialoglycans serve as ligands for self-Siglec receptors, functioning as self-associated molecular patterns (SAMPs) that maintain macrophage in an immunosuppressive state (**Fig. 4d**) ^55, 56^. To investigate whether engineered bacteria-mediated desialylation could reverse the immunosuppression, we pre-treated the THP-1-derived macrophage with either the sialidase-active strain EcN^Δ^*^pgi^*-SS or sialidase-inactive EcN^Δ^*^pgi^*-S^E647A^S strain. These pre-treated THP-1-derived macrophages were then co-cultured with WT SK-BR-3 tumor cells at an effector-to-target ratio of 5:1 for 8 hours. Notably, macrophages exposed to EcN^Δ^*^pgi^*-SS demonstrated significantly enhanced tumor cell phagocytosis compared to those treated with EcN^Δ^*^pgi^*-S^E647A^S strain (**Fig. 4e**). This indicates that bacterial desialylation of macrophages alleviates self-inhibition and potentiates their tumor-phagocytic capacity.

### Engineered bacteria synergize with PD-L1 antibody to suppress mouse subcutaneous tumor growth

Given that engineered bacteria exhibited potency to enhance the anti-tumor activity of immune cells, we subsequently sought to assess the immunotherapeutic efficacy of engineered bacteria in vivo. Poorly immunogenic “cold” 4T1 tumors typically exhibit limited responsiveness to immune checkpoint inhibitors ^57, 58^. Therefore, we evaluated the anti-tumor efficacy of combining engineered bacteria with an anti-PD-L1 antibody (αPD-L1) on a mouse model bearing 4T1-HER2 tumors. Based on dose-titration experiments, a suboptimal dose of αPD-L1 (50 μg per mouse), which demonstrates minimal single-agent activity, was selected for the combination study (**Supplementary Fig. 13)**. Mice bearing 4T1-HER2 tumors were assigned to one of eight treatment groups: PBS (G1), αPD-L1 alone (G2), or αPD-L1 combined with various bacterial strains, including EcN^Δ^*^pgi^* (G3), EcN^Δ^*^pgi^*-S (G4), EcN^Δ^*^pgi^*-SA (G5), EcN^Δ^*^pgi^*-SS (G6), EcN^Δ^*^pgi^*-S^E647A^AS (G7), and EcN^Δ^*^pgi^*-SAS (G8). The αPD-L1 (50 μg in 100 μl PBS) was administrated intraperitoneally, while the engineered bacteria (1 × 10^5^ CFU in 100 μl PBS) were delivered intravenously via the tail vein. Both agents were administered every three days for a total of five doses (**Fig. 5a**).

**Figure 5.**
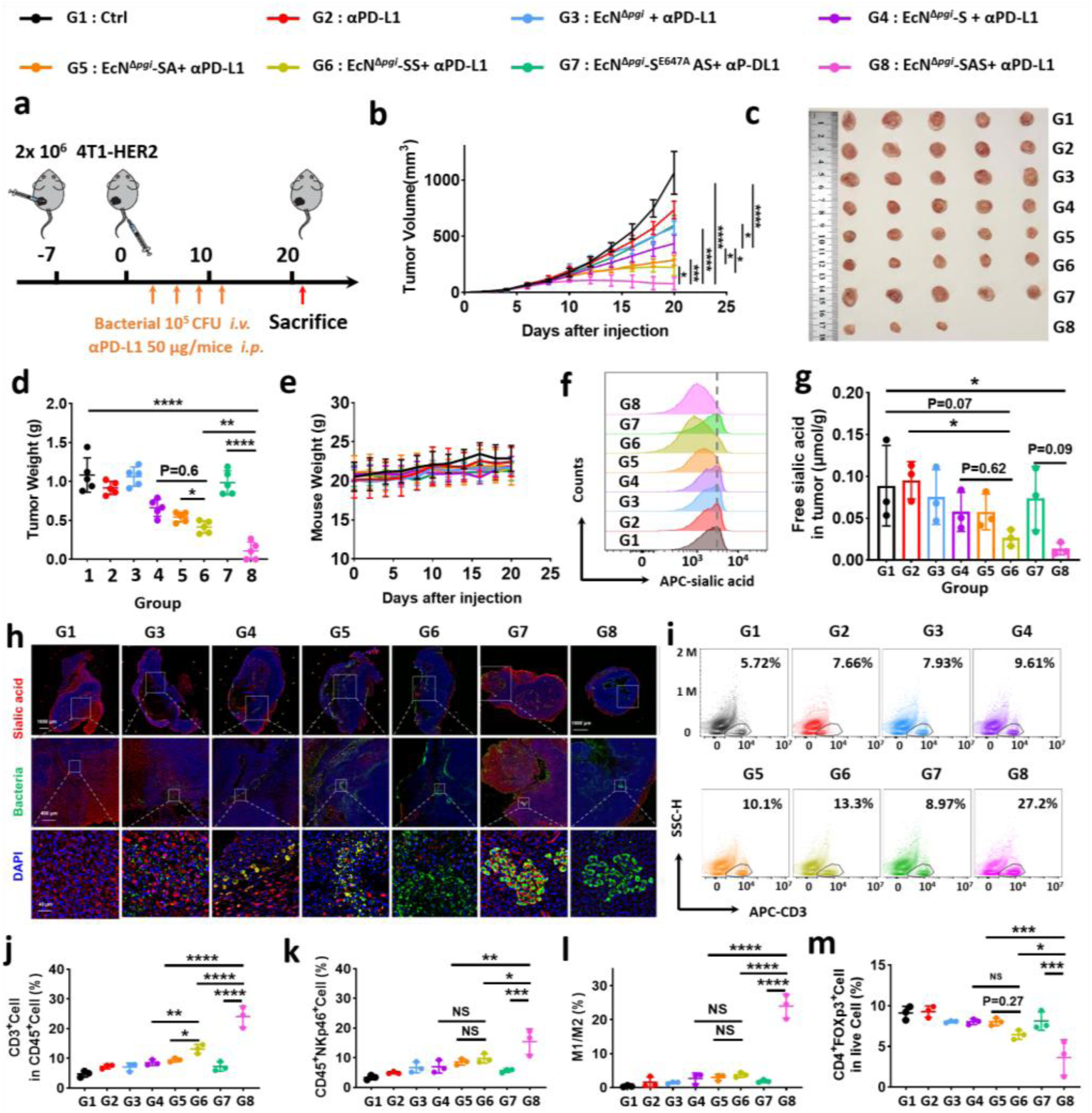
Engineered bacteria synergize with anti-PD-L1 antibody to suppress subcutaneous tumors. **a**, Schematic of the combinational therapeutic regimen for inhibiting 4T1-HER2 subcutaneous tumor growth. Tumor-bearing mice received intravenous administration of the indicated bacteria (10^5^ CFU per mouse) and intraperitoneal administration of anti-PD-L1 antibody (50 μg per mouse) on days 3, 6, 9, 12 and 15. **b**, Tumor growth curves of 4T1-HER2 tumor-bearing mice receiving the indicated bacteria treatments combined with anti-PD-L1 antibody. **c**, Representative images of excised tumors from mice in each treatment group. **d,** Quantification of tumor weights across treatment groups. **e**, Longitudinal monitoring of mouse body weights throughout the treatment period. **b, d**, **e,** Data are shown for n = 5 biologically independent mice per group. Statistical analysis was performed using one-way ANOVA followed by Tukey’s multiple comparisons test. \**P* < 0.05, \*\**P* < 0.01, \*\*\**P* < 0.001 and \*\*\*\**P* < 0.0001. **f,** Flow cytometry profiles of CD45⁻ tumor cells stained with Sambucus nigra agglutinin (SNA-Cy5) across treatment groups. **g**, Quantification of free sialic acid concentrations in tumor tissue supernatants measured via LC-MS. Data are shown for n = 3 biologically independent mice per group. Statistical analysis was performed using one-way ANOVA followed by Tukey’s multiple comparisons test. **h,** Immunofluorescence staining of tumor sections from mice across treatment groups. Red: sialic acid; green: bacteria; blue: nuclei. **i,** Flow cytometry profiles of tumor-infiltrating CD3⁺ T cells. Quantification of tumor-infiltrating CD3^+^ T cells (**j**), NK cells (**k**), M1/M2 macrophage ratio (**l**) and Foxp3^+^ Treg cells (**m**). **g, j, k, l, m,** Data are shown for n = 3 biologically independent mice per group. Statistical analysis was performed using one-way ANOVA followed by Tukey’s multiple comparisons test. \**P* < 0.05, \*\**P* < 0.01, \*\*\**P* < 0.001 and \*\*\*\**P* < 0.0001.

Monotherapy with αPD-L1 marginally inhibited tumor growth by 32% (G2). Co-administration of EcN^Δ^*^pgi^* did not enhance anti-tumor efficacy compared to αPD-L1 alone (G3). In contrast, combining αPD-L1 with the sialidase-active engineered bacterium EcN^Δ^*^pgi^*-S significantly enhanced anti-tumor activity, inhibiting tumor growth by 60% (G4). This potentiation was further augmented by co-administration with either EcN^Δ^*^pgi^*-SA (equipped with an anti-HER2 affibody, G5) or EcN^Δ^*^pgi^*-SS (harboring the sialic acid catabolism gene cluster, G6). Remarkably, combination of αPD-L1 and EcN^Δ^*^pgi^*-SAS (G8) almost completely suppressed tumor growth. Conversely, the sialidase-inactive mutant EcN^Δ^*^pgi^*-S^E647A^AS (G7) failed to enhance anti-tumor efficacy. Moreover, mice body weights remained stable across all treatment groups, confirming the biosafety of these combinational treatments. (**Fig. 5e**). Collectively, the results demonstrate that sialidase activity, anti-HER2 targeting, and sialic acid catabolism synergistically optimize engineered bacteria to potentiate αPD-L1 therapy. After mice sacrifice, tumors were harvested, processed into single-cell suspensions, and surface sialylation on CD45⁻ tumor cells were quantified via SNA-Cy5 staining. Tumor cells from G5, G6 and G8 groups exhibited significantly reduced surface sialylation. The lowest levels were observed in G6 and G8, both groups harboring engineered bacteria with active sialidase and the sialic acid catabolism gene cluster (**Fig. 5f**). These data indicated that the catabolism gene cluster potentiates engineered bacteria-mediated tumor cell desialylation. Furthermore, free sialic acid levels in tumor suspension supernatants were also quantified by LC-MS. Consistent with desialylation results, G6 and G8 group showed minimal free sialic acids accumulation. In contrast, G4 and G5 group exhibited significantly elevated free sialic acids concentrations (**Fig. 5g**). These results demonstrate that while active sialidase liberates sialic acids, the bacterial catabolic machinery in G6 and G8 groups effectively scavenges them.

Given that all engineered bacteria harbored an sfGFP reporter, engineered bacteria colonization and tumor sialylation levels were assessed via immunofluorescence. Tumor sections co-stained with SNA-Cy5 and DAPI revealed spatial relationships between bacterial localization and tumor cell desialylation. Robust green fluorescence confirmed successful bacterial colonization across groups G3-G8. Intriguingly, strains in G4-G8 groups exhibited enhanced tumor targeting and adhesion relative to the strain in G3 group, suggesting that either bacterial surface-exposed sialidase or anti-HER2 affibody enhanced tumor cell adhesion. Significantly higher sialylation was observed in the sialidase-inactive control group (G7) compared to G8 group, confirming that active sialidase is essential for tumor desialylation. Notably, pericellular sialic acid levels at bacteria-adhered tumor cells were markedly lower in G6 and G8 than in G4 and G5, further validating that the catabolic gene cluster enhances tumor desialylation efficacy by scavenging liberated sialic acid (**Fig. 5h**).

Tumor desialylation of tumor implied TME remodeling. Therefore, tumor infiltrating immune cells were subsequently examined. Consistent with the desialylation levels, flow cytometry analysis revealed that G8 treatment induced maximal intratumoral CD3⁺ T cell infiltration (27.2% of live CD45^+^ cells), followed by the G6 group (13.3% of live CD45^+^ cells) (**Fig. 5i, j**). Furthermore, the G8 group exhibited the most significant elevation in NK cells infiltration and M1/M2 m macrophage ratio, alongside the greatest reduction in CD4⁺FoxP3⁺ regulatory T cells (**Fig. 5k-m and Supplementary Fig. 14-18)**. Immunofluorescence analysis also corroborated these data, demonstrating enhanced infiltration of CD8⁺ T cells, NK cells, and M1 macrophages in G8-treated tumors compared to other groups, confirming that combinatorial therapy heightened immunocyte recruitment and reprogrammed the immunosuppressive microenvironment **(Supplementary Figs. 19-21)**. Additionally, serum levels of IFN-γ, TNF-α and IL-12p70 were markedly elevated in G8-treated mice, indicating systemic immunostimulation **(Supplementary Fig. 22)**. Histological assessment of mouse tissues detected no abnormalities across treatment groups **(Supplementary Fig. 23)**. Collectively, these results establish that engineered EcN^Δpgi^-SAS synergizes with PD-L1 blockade via tumor-directed desialylation, metabolic scavenging, and immunopotentiation to remodel the tumor immune microenvironment and suppress tumor growth.

### Engineered bacteria inhibit tumor growth in an AOM/DSS-induced mouse colorectal cancer model

The oral administration of probiotics represents a preferred, effective alternative to intravenous delivery, offering sustained mitigation of systemic toxicities. Our previous studies demonstrated that oral administration of 10^9^ CFU engineered EcN could persist in the murine intestine for over 6 days, suggesting that the potential for long-term therapeutic effects within the gut ^41^. Given the established capacity of engineered bacteria to remodel the tumor immune microenvironment with high efficiency, we subsequently evaluated their therapeutic efficacy in an AOM/DSS-induced mouse colorectal cancer (CRC). Briefly, mouse CRC tumors were induced in 3-week-old C57BL/6c mice over 10 weeks using a single azoxymethane (AOM) injection followed by five cycles of 3% dextran sulfate sodium (DSS) in drinking water, with weekly alternation to water-only intervals to mitigate DSS-associated severe colitis and mortality (**Fig. 6a and Supplementary Fig. 24)**. In the eighth week, the CRC-bearing mice were randomly assigned to five groups. During the following three weeks, each group received various treatments, including PBS (G2), the sialidase-inactive strain EcN^Δ^*^pgi^*-S^E647A^ (G3), sialidase-active strain EcN^Δ^*^pgi^*-S (G4), sialidase-active with anti-HER2 affibody strain EcN^Δ^*^pgi^*-SA (G5) and EcN^Δ^*^pgi^*-SAS (G6). Typically, 100 μL of a bacterial suspension in PBS at a concentration of 1 × 10^10^ CFU mL^−1^ was administered via oral gavage every three days for a total of five doses. Mice that did not receive AOM/DSS treatment served as the healthy control group (G1). At week 11, the mice were euthanized and colons were measured due to the established inverse correlation between colon length and colonic lesion severity. Mice in G2 and G3 groups had the shortest colon length and displayed visible intestinal tumors. In contrast, sialidase-active strain-treated groups (G4 and G5) showed significantly longer colons than G2 and G3, while EcN^Δpgi^-SAS administration (G6) resulted in colon lengths comparable to the control group (G1). Importantly, G6 mice developed substantially fewer tumors than all other treatment groups. These results demonstrate that sialidase-expressing engineered bacteria attenuate colorectal tumorigenesis in mice, an effect significantly augmented by sialic acid catabolic gene clusters (**Fig. 6b, c**). Consistently, immunofluorescence staining images revealed the lowest Ki-67 expression levels in colonic sections of G6 mice compared to other treatment groups, indicative of suppressed tumor-cell proliferation in colonic tissues **(Supplementary Fig. 25)**. Histological analysis via H&E staining demonstrated severe tissue damage, crypt destruction, and tumor infiltration in mice across G2-G5 groups. In contrast, G6 group mice exhibited markedly preserved mucosal architecture with intact crypt integrity comparable to the control group, indicating near-normal histology (**Fig. 6d-e**).

**Figure 6.**
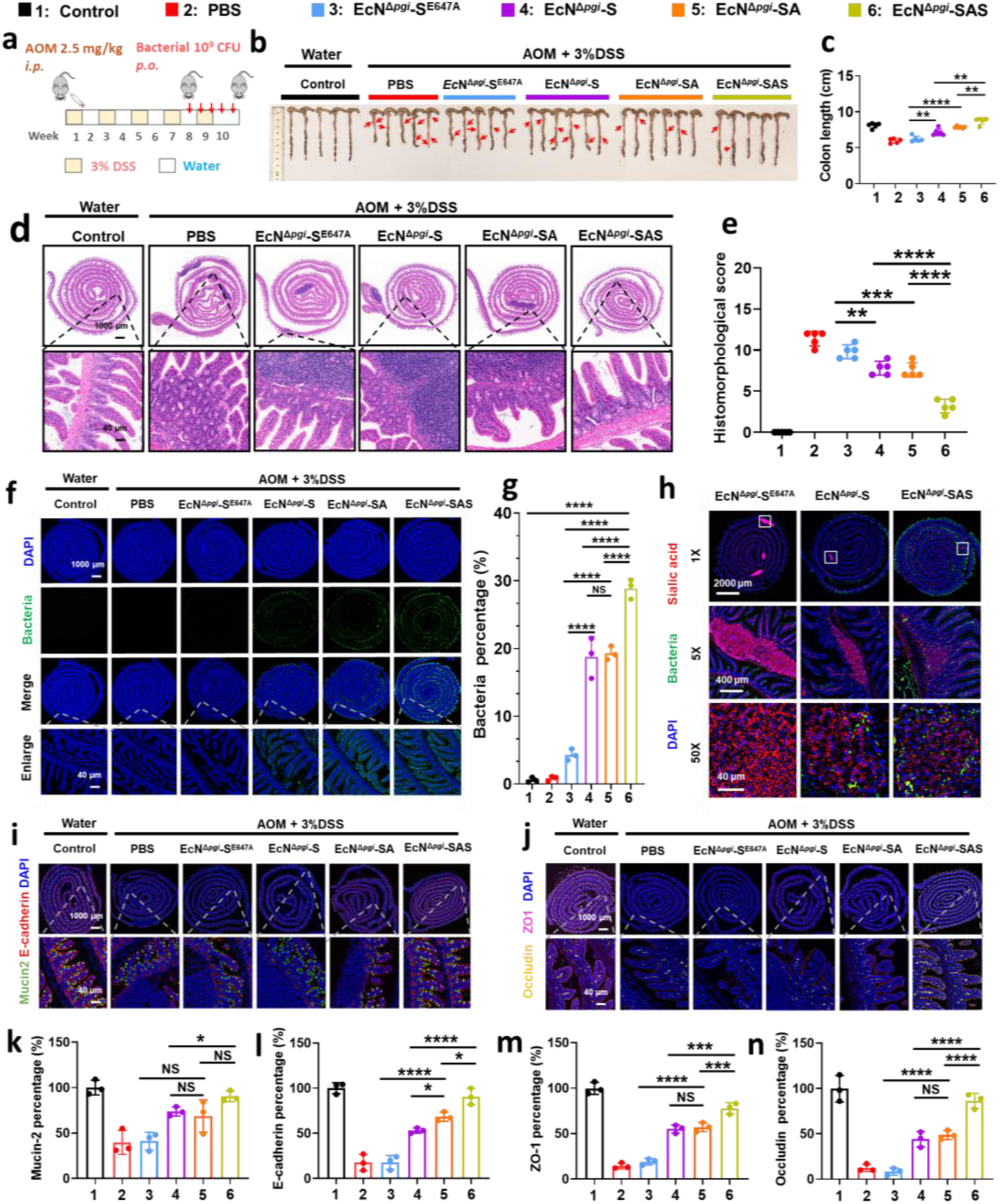
Engineered bacteria suppresses AOM/DSS-induced colorectal cancer and restore intestinal barrier integrity. **a**, Schematic of AOM/DSS-induced colorectal cancer model and experimental procedure. Mice were intraperitoneally injected with AOM (10 mg kg⁻¹). One week later, 5 cycles of 3% (w/v) DSS (7 d DSS/7 d water) were applied to establish the AOM/DSS colorectal cancer model. Tumor-bearing mice received oral administration of 1 × 10⁹ CFU per mouse of indicated bacterial strains every 3 d for five doses. **b**, **c**, Representative images (**b**) and length (**c**) of mice colons across treatment groups. Data are shown for n = 5 biologically independent mice per group. **d, e,** Representative H&E-stained sections (**d**) and colon damage scores (**e**) of colon tissues from each treatment group. Data are shown for n = 5 biologically independent mice per group. **f, g**, Representative immunofluorescence staining images (**f**) and semi-quantitative analysis (**g**) of bacteria colonization in mice colon tissue sections after treatment. Green: bacteria; blue: nuclei. Data are shown for n = 3 independent experiments. **h**, Representative immunofluorescence staining images of bacteria and tumor sialylation in mice colon tissue sections after treatment. Red: sialic acid; green: bacteria; blue: nuclei. **i-n**, Representative immunofluorescence staining images (**i, j**) and semi-quantification of adhesion molecules mucin 2 (**k**) and E-cadherin (**l**), tight junction proteins ZO-1 (**m**) and occludin (**n**). Blue: nuclei; red: E-cadherin; green: mucin 2; pink: ZO1; yellow: occludin. Data are presented as mean ± s.d. from n = 3 independent experiments. **c, e, g, k, l, m, n,** Statistical analysis was performed using one-way ANOVA followed by Tukey’s multiple comparisons test. \**P* < 0.05, \*\**P* < 0.01, \*\*\**P* < 0.001 and \*\*\*\**P* < 0.0001. NS, not significant

Since all engineered bacteria strains encoded a chromosomally integrated sfGFP reporter gene, colonization and distribution of engineered bacteria in mice intestinal tract could be visualized. Immunofluorescence staining images of colonic sections revealed negligible bacterial colonization in G3 group mice, whereas substantial sfGFP signals were observed in G4-G6 groups (**Fig. 6f**). Notably, mice in G6 group exhibited the most robust bacterial epithelial colonization, with sfGFP fluorescence intensity significantly elevated by 9% compared to both G4 and G5 group, reflecting enhanced bacterial epithelia adherence. This enhanced colonization is probably attributable to the sialic acid catabolism system in EcN^Δ^*^pgi^*-SAS, where the cleaved tumor-derived sialic acids serve as a carbon source. This metabolic adaptation alleviates growth suppression and supports bacterial proliferation within the sialic acid-rich TME, thereby facilitating mucosal colonization (**Fig. 6g**). Furthermore, tumor sections sialylation was also analyzed via SNA-Cy5 and DAPI staining. Immunofluorescence imaging revealed obvious increased bacterial adhesion to tumor cells and reduced sialylation signals in group G4 relative to G3, demonstrating engineered bacteria-mediated desialylation. Crucially, G6 group mice exhibited the most pronounced sialylation reduction among treated groups, confirming that the sialic acid catabolic system facilitates in-situ cleavage and assimilation of tumor-derived sialic acids (**Fig. 6h and Supplementary Fig. 26)**.

To further evaluate intestinal structure and barrier integrity, the epithelial cells of mouse colonic tissues were examined. Specifically, Mucin 2, a core glycoprotein of the mucus layer, and E-cadherin, a key mediator for cell-cell adhesion and epithelial integrity, were analyzed by immunofluorescence staining (**Fig. 6i, 6j**). Immunofluorescence results showed modestly increased levels of Mucin 2 and E-cadherin in the colonic tissue of G4 and G5 groups. Notably, the EcN^Δ^*^pgi^*-SAS-treated G6 group exhibited a remarkably elevated levels of Mucin 2 and E-cadherin, reaching 90% and 91% of the expression levels in healthy control mice, respectively. In contrast, negligible change was observed in the sialidase-inactive strain group (G3) (**Fig. 6k, 6l**). Similarly, the levels of two pivotal tight junction proteins, zonula occludens 1 (ZO1) and occludin, which are indispensable for maintaining intestinal epithelial barrier, were substantially increased in the mouse groups treated with sialidase-active strains. Typically, the G6 group displayed the highest levels of ZO1 and occluding, which were with 78% and 86% of healthy control levels, respectively (**Fig. 6m, 6n**). Collectively, these results indicate that engineered bacteria EcN^Δ^*^pgi^*-SAS not only inhibits tumor cell proliferation but also efficiently restores impaired colonic epithelial barrier function in CRC-bearing mice.

## Discussion

Hypersialylation constitutes a hallmark feature of the immunosuppressive TME, driving tumor immune evasion, metastasis, and resistance to immunotherapy. Antibody-sialidase conjugates that selectively remove tumor sialylation enhance antitumor immunity and suppress tumor progression in vivo ^22, 24, 25^. Similarly, a bispecific T-cell engager (BiTE)-sialidase fusion protein was recently shown to augment tumor susceptibility to BiTE-mediated cytolysis through targeted desialylation^23^. However, such sialidase-conjugates exhibit intrinsic limitations including nonspecific desialylation and systemic instability. Moreover, tumors can potentially reuse liberated sialic acid to restore hypersialylation. To address these challenges, we engineered the probiotic EcN strain EcN^Δ^*^pgi^*-SAS incorporating multifunctional modules, including HER2-directed tumor anchoring, hypoxia-inducible sialidase expression, autonomous sialic acid catabolism, and tumor-restricted conditional proliferation. The hypoxia-responsive surface-displayed sialidase enables in situ cleavage of tumor-associated sialic acid chains. The liberated sialic acid is then irreversibly catabolized by the integrated sialic acid scavenging system through competitive uptake and conversion into bacterial glycolytic intermediates. This sequential action prevents both nonspecific desialylation side effects and liberation-driven sialic acid reincorporation by tumors. Furthermore, we rewired bacterial carbon metabolism via disruption the *pgi* gene to abolish glucose utilization, forcing metabolic dependence on sialic acid. EcN^Δpgi^-SAS thus exclusively proliferates in sialic acid-rich tumor niches by scavenging and catabolizing TME-derived sialic acid. This metabolism shift enhances biosafety, eliminating uncontrolled systemic growth, reducing biofilm formation, and accelerating off-tumor bacterial clearance. EcN^Δpgi^-SAS thus exhibits minimal systemic colonization while maintaining robust tumor accumulation, improving host survival and optimizing therapeutic safety for clinical translation.

In vitro and in vivo analyses confirm that engineered bacteria remodel the immunosuppressive TME, counteract sialylation-mediated immune evasion, and synergize with PD-L1 blockade to suppress subcutaneous tumor growth in mice. Critically, monotherapy with this strain potently inhibits AOM/DSS-induced mouse colorectal tumor growth and restores impaired colonic barrier function. During manuscript preparation, a contemporaneous study reported an engineered EcN co-expressing a *Salmonella typhimurium* sialidase and a photosensitizer for photodynamic therapy ^59^. However, this system cannot catabolize liberated sialic acid and depends on light-driven photodynamic activation, which suffers from poor tissue penetration and is inapplicable to intestinal tumors. Our approach thus provides a targeted, systemically safe, and clinically viable microbial immunotherapy strategy.

Despite its efficacy, this strategy has limitations requiring optimization prior to clinical translation. HER2-specific targeting currently restricts application to HER2^+^ tumors. Incorporation of multi-biomarker recognition modules could expand applicability. Furthermore, while the engineered bacteria suppress AOM/DSS-induced colitis-associated colorectal cancer, its impact on endogenous gut microbiota requires further evaluation.

## Methods

### Bacterial strains and culture conditions

*Escherichia coli* Nissle 1917 (EcN) was used as the parental strain for all genetic engineering. Bacteria were routinely cultured in Luria-Bertani (LB) broth or on LB agar at 37°C. All bacteria were grown with appropriate antibiotics selection (100 μg ml^-1^ of ampicillin, 50 μg ml^-1^ of kanamycin and 34 μg ml^-1^ of chloramphenicol). Anaerobic cultures were maintained in a hypoxia workstation with a gas mixture of 88% N2, 10% CO2, and 2% O2. Bacterial growth was monitored via OD600 measurements using a multi-mode microplate reader (BioTek Instruments).

### Construction of plasmids and genetic circuits

Codon-optimized sialidase A gene (*nanA*) from *Streptococcus pneumoniae* was synthesized by Genewiz and sub-cloned into the pET47b plasmid using restriction sites of EcoRI and Sall, yielding pET47b-*Sp*NanA. Plasmid of inactive mutant *Sp*NanA^E647A^ was constructed via site-directed mutagenesis using the wild-type pET47b-*Sp*NanA plasmid as DNA template. The constructed plasmids were transformed into BL21(DE3) cells for expression and purification. For bacterial surface display, *nanA* was fused to the Lpp-OmpA anchoring domain coding sequence under the control of the hypoxia-responsive *fnrS* promoter (P*fnrS*). Concurrently, an anti-Her2 affibody coding sequence was fused to the AIDA-I autotransporter coding sequence, driven by the constitutive *lacUV5* promoter (P*lacUV5*). These expression cassettes were assembled into the previously reported stable plasmid pSYSU-1 ^41^ using a One Step Cloning Kit (Vazyme Biotech). The sialic acid catabolic gene cluster, including *nanT* (encoding the sialic acid transporter), *nanA* (encoding the N-acetylneuraminate lyase), *nanK* (encoding the N-acetylmannosamine kinase), *nanE* (encoding the N-acetylmannosamine-6-phosphate 2-epimerase), *nagA* (encoding the N-acetylglucosamine-6-phosphate deacetylase), and *nagB* (encoding the glucosamine-6-phosphate deaminase), was codon-optimized and synthesized by Genewiz. Each gene was C-terminally tagged for detection as follows: *nanT* with a T7-tag, *nanA* with an HA-tag, *nanK* with a MYC-tag, *nanE* with a V5-tag, *nagA* with a FLAG-tag, and *nagB* with an S-tag. The tagged cluster was cloned into the downstream of the P*lacUV5* promoter, with *nanR* incorporated as a sialic acid-responsive repressor. This assembly was integrated into a stable plasmid pSYSU-2 containing a Hok/Sok stabilization system. All cloned sequences were verified by Sanger sequencing (Sangon Biotech). Plasmids were electroporated (25 μF, 200 Ω, 1.8 kV) into EcN using a MicroPulser system (Bio-Rad Laboratories). All primers used in this study are listed in **Supplementary Table 1.**

### Construction of attenuated EcN mutant strains

For attenuated EcN generation, the *pgi* gene encoding phosphoglucose isomerase in EcN was deleted using a pEcCas/pEcgRNA system as previously described ^60^. A 20-nt guide RNA targeting the start codon region of *pgi* was cloned into the pEcgRNA plasmid. Homology arms flanking *pgi* gene (500 bp each) were amplified from EcN genomic DNA and assembled into pEcgRNA via Gibson assembly. The resulting plasmid was co-transformed with pEcCas9 into EcN. Transformants were selected on LB agar supplemented with kanamycin (50 μg mL⁻¹), spectinomycin (50 μg mL⁻¹), and L-arabinose (0.5%, w/v). Verification of *pgi* deletion was performed through colony PCR using primers spanning the gene locus, followed by Sanger sequencing.

### Sialidase expression, purification, and in vitro activity assay

Recombinant *Sp*NanA and its inactive mutant *Sp*NanA^E647A^ were expressed in *E. coli* BL21 (DE3) with an N-terminal His-tag. Briefly, transformed BL21 (DE3) cells were cultured at 37 °C until reaching an OD600 of 0.6–0.8, induced with 0.5 mM isopropyl β-D-1-thiogalactopyranoside (IPTG), and further incubated overnight at 16 °C. Cells were harvested by centrifugation (5,000 × g, 15 min, 4°C), resuspended in lysis buffer (20 mM Tris-HCl, 300 mM NaCl, 10 mM imidazole, 1 mM PMSF, pH 8.0), and lysed via high-pressure homogenization. Following centrifugation (20,000 × g, 30 min, 4°C), soluble fractions were applied to a Ni^2+^-NTA affinity column. After sequential washing steps with wash buffer (20 mM imidazole in lysis buffer), bound proteins were eluted with elution buffer (500 mM imidazole, 20 mM Tris-HCl, 300 mM NaCl, pH 8.0) and further purified through size-exclusion chromatography using an ÄKTA pure FPLC system (Cytiva) with PBS (pH 7.4) mobile phase. Protein concentrations were determined using a BCA protein assay kit (Thermo Fisher Scientific). Purified enzymes were incubated with fetuin (Sigma-Aldrich) at a 1:100 enzyme-to-substrate molar ratio in PBS buffer (pH 7.4) for 1 h at 37 °C. Reactions were terminated by boiling in SDS-PAGE loading buffer for 10 min, followed by analysis via 10% SDS-PAGE with Coomassie Brilliant Blue staining. For MALDI-TOF MS analysis, fetuin samples were desalted using C18 ZipTips (Millipore) and analyzed in linear positive ion mode on a Bruker UltrafleXtreme MALDI-TOF mass spectrometer.

### Western blot analysis of target protein expression in EcN-SAS strain

Expression of the bacterial surface-displayed *Sp*NanA and anti-HER2 affibody was assessed by culturing bacteria in LB medium under normoxic or hypoxic conditions until OD600 value reached 0.8. For the detection the expression of sialic acid catabolic genes, bacteria in log phase were incubated with a gradient concentrations of sialic acid ranging from 0-1000 μg/mL for 2 h. The bacteria cultures were harvested by centrifugation at 5,000 × g for 10 min at 4°C, and lysed via sonication (30 W output, 3 s pulse-on, 3 s pulse-off) on ice in RIPA buffer supplemented with protease inhibitor cocktail (Roche). Protein concentrations were quantified using a BCA assay. Equivalent samples (30 μg protein per lane) were resolved by 12% SDS-PAGE andelectrotransferred to PVDF membranes (Millipore). The membranes were blocked for 2 h at ambient temperature with 3% (w/v) BSA in TBST, followed by overnight incubation at 4°C with the indicated primary antibodies against epitope tags: anti-HA for *Sp*NanA, anti-Myc for anti-HER2 affibody, anti-T7 for NanT, anti-HA for NanA, anti-Myc for NanK, anti-V5 for NanE, anti-FLAG for NagA, and anti-S-tag for NagB. After washing, membranes were incubated with HRP-conjugated secondary antibodies (1:10000) for 2 h at 25°C. Bacterial GAPDH was used as an internal reference. Protein signals were detected using enhanced chemiluminescence substrate (NCM Biotech) and imaged with a GeneGnome XRQ chemiluminescence imaging system (Syngene).

### Quantitative PCR analysis of *nanT* transcription in EcN-SAS strain

Total RNA was extracted from bacterial cultures using TRIzol reagent (Invitrogen) according to the manufacturer’s protocol. Extracted RNA concentration was quantified using a NanoDrop 2000 spectrophotometer (Thermo Fisher Scientific). Complementary DNA (cDNA) was synthesized from 1 μg of total RNA with GoScript Reverse Transcriptase (Promega) and random hexamer primers. Quantitative real-time PCR (qPCR) was performed using GoTaq qPCR Master Mix (Promega) on a StepOnePlus Real-Time PCR System (Thermo Fisher Scientific) with the following cycling parameters: 95°C for 10 min, then 40 cycles of 95°C for 15 s and 60°C for 1 min. Transcription of *nanT* was normalized to 16S ribosomal RNA and analyzed using the ΔΔCt method.

### LC-MS analysis of sialic acid catabolism in EcN-SAS strain

The bacteria were cultured in LB medium supplemented with 2 mM sialic acid until OD₆₀₀ reached 0.5. At indicated time points (0, 30, 60, 120, 240 min), culture supernatants were collected by centrifugation (13,000 × g, 5 min), filtered through 0.22 μm membranes, and analyzed for residual sialic acid content by LC-MS. For intracellular metabolite profiling, bacterial pellets were washed twice with ice-cold PBS, lysed by freeze-thaw cycles in methanol/water (80:20, v/v), and processed for LC-MS quantification for sialic acid and catabolic intermediates, including N-acetylmannosamine (ManNAc), N-acetyl-mannosamine-6-phosphate (ManNAc-6-P), N-acetylglucosamine-6-phosphate (GlcNAc-6-P), D-glucosamine-6-phosphate (GlcN-6-P) and fructose-6-phosphate (Fru-6-P). LC-MS analysis was performed on a Q Exactive HF-X Orbitrap mass spectrometer coupled to a Dionex UltiMate 3000 UHPLC system (Thermo Fisher Scientific) in SIM mode. Metabolites separation was achieved on a InfinityLab Poroshell 120 EC-C18 column (3×150 mm, 1.9-Micron) with a mobile phase of acetonitrile and 20 mM ammonium acetate (pH 9.0).

### Mammalian cell culture

The 4T1, human HER2-transfected 4T1 cells (4T1-HER2), MDA-MB-231, SK-BR-3, Hela, MCF-7, CT26, A549, AGS and MC38 cells were obtained from ATCC. Cells were cultured in either RPMI-1640 or DMEM supplemented with 10% FBS, 100 U ml^-1^ penicillin, and 100 μg ml^-1^ streptomycin. Cell cultures were maintained at 37°C in a humidified 5% CO2 atmosphere. HER2 overexpression in 4T1-HER2 was validated by flow cytometry using an anti-HER2 antibody (BioLegend). Routine mycoplasma screening was performed using the MycoAlert PLUS Mycoplasma Detection Kit (Lonza).

### Flow cytometry of bacteria-mediated tumor cell desialylation

Tumor cells were seeded in 24-well plates at 2 × 10^5^ cells per well and allowed to adhere overnight. Engineered bacteria were added at indicated cell-to-bacteria ratios (1:10, 1:50 and 1:100) under normoxic or hypoxic conditions for 4 h. Post co-culture, tumor cells were extensively washed with PBS, detached via trypsin-EDTA treatment, and stained with Cy5-conjugated *Sambucus nigra* agglutinin (SNA-Cy5, Vector Laboratories) at a 1:200 dilutions for 30 min at 4°C. Flow cytometry was performed on a CytoFLEX instrument (Beckman Coulter), with data analyzed using FlowJo software (v10.8.1). For kinetic profiling, tumor-bacteria (cell-to-bacteria ratio of 1:10) co-culture were sampled at indicated intervals (0, 30, 60, 90, 120, 150, 180, 210 min).

### Confocal microscopy of bacteria-mediated tumor cell desialylation

Tumor cells were seeded onto glass coverslips placed in 24-well culture plates. The tumor cells were then co-cultured for 4 h with bacteria bearing a genomically integrated sfGFP reporter as described above. After co-culture, cell samples were fixed with 4% paraformaldehyde for 15 min at room temperature, blocked with 1% BSA in PBS for 1 h at room temperature, and stained with SNA-Cy5 (1:200, Vector Laboratories) to label sialic acid for 30 min at room temperature. Nuclei were counterstained with Hoechst 33342 for 5 min at room temperature. Coverslips were imaged on a Zeiss LSM 880 confocal microscope with a 63× oil immersion objective. Images were processed using ZEN software (Zeiss).

### 13C-labeled sialic acid competition between tumor cells and bacteria

Tumor cells were cultured for 72 h in medium supplemented with 2 mM ^13^C-labeled sialic acid (Cambridge Isotope Laboratories) to metabolically label surface glycans. Labelled cells were washed extensively with PBS and co-cultured with bacteria under hypoxic conditions for 12 h. After co-culture, cells were washed three times with PBS to detach loosely adherent bacteria. Harvested bacteria were centrifuged at 4,000 × g for 5 min to form bacterial pellet, followed by three additional washes with PBS. Bacterial pellets were resuspended in PBS, lysed by sonication, and immediately mixed with an equal volume of ice-cold methanol for protein precipitation. The resulting supernatant was collected, and intracellular levels of ^13^C-sialic acid and ^13^C-ManNAc-6-P were analyzed via LC-MS in selected ion monitoring (SIM) mode. The remaining adherent tumor cells were collected by Trypsin-EDTA treatment, followed by centrifugation at 500 × g for 5 min. The collected cells were washed three times with PBS and incubated with 500 μM purified *Sp*NanA for 1 h at 37°C to cleave residual surface sialic acids. The supernatant containing cleaved ^13^C-sialic acid was collected and analyzed by LC-MS.

### Growth of attenuated bacteria with different carbon sources

Overnight cultures of attenuated bacteria were diluted 1:100 in fresh LB medium with or without different supplements (glucose at 5, 10, and 20 mM; fructose-6-phosphate at 5 mM; sialic acid at 5 mM). Two hundred microliters of each diluted culture were transferred to 96-well plates, then incubated at 37 °C under continuous shaking in a BioTek microplate reader. Optical density at 600 nm (OD600) was recorded every 15 min for 500 min.

### Biofilm formation assay

Bacterial strains were grown overnight in LB medium, diluted to an OD600 of 0.5, and transferred to 24-well polystyrene plates (1 mL per well). After 24 h of static incubation at 37 °C to allow biofilm formation, planktonic bacteria were removed by gentle washing with PBS. Attached biofilms were fixed with methanol for 15 min, stained with 0.1% (w/v) crystal violet for 20 min, washed repeatedly with PBS, and solubilized in 33% (v/v) acetic acid. The absorbance of the solubilized crystal violet was measured at 595 nm. All assays were performed with six biological replicates per strain.

### Sensitivity of bacteria to blood components

Fresh human whole blood was obtained from healthy volunteer donors at the First Affiliated Hospital of Sun Yat-sen University. This study was approved by the Clinical Research Ethics Committee, the First Affiliated Hospital of Sun Yat-sen University, and written informed consent was acquired from all participating donors. All procedures complied with the Declaration of Helsinki. Harvested bacteria were washed twice with PBS and adjusted to a concentration of 1 × 10⁶ CFU mL⁻¹ in either PBS or 90% (v/v) whole blood (diluted in PBS). Samples were incubated at 37 °C with gentle rotation. At 0, 6, and 12 h of incubation, aliquots were serially diluted and plated on LB agar for CFU enumeration. Bacterial survival was calculated as the percentage of viable bacteria relative to the initial count at time zero. Harvested bacteria were washed twice with PBS and adjusted to a concentration of 1 × 10⁶ CFU ml⁻¹ in either PBS or 90% (v/v) whole blood (diluted in PBS). Samples were incubated at 37 °C with gentle rotation. At 0, 6, 12 h of incubation, aliquots were serially diluted and plated on LB agar for CFU enumeration. Bacterial survival was calculated as the percentage of viable bacteria relative to the initial count at time zero.

### Mouse survival and bacterial organ colonization

The animal experiment procedures were approved by the Institutional Animal Care and Use Committee of Sun Yat-sen University (approval number: 2025003715) and performed in accordance with the ethical guidelines of Sun Yat-sen University. All the experiments were conducted in the Laboratory Animal Center of Sun Yat-sen University. Female BALB/c and C57BL/6 mice (6–8 weeks old) were obtained from Guangzhou Laboratory Animal Center and housed under specific pathogen-free conditions. For survival studies, 2 × 10^6^ MC38 cells were injected subcutaneously into the right flank of C57BL/6 mice. Tumor volumes were measured every 2-3 days with digital calipers and calculated as (length × width²)/2. When tumors reached a volume of 50-100 mm³, mice were randomly assigned to different treatment groups. MC38 tumor-bearing C57BL/6 mice (n = 10 mice per group) were injected intravenously with a high dose of 1 × 10⁷ CFU of EcN-SAS or attenuated EcN^Δ^*^pgi^*-SAS every two days for a total of five doses. Mice were monitored daily for up to 30 days, and survival curves were compared using the log-rank test. For bacterial organ colonization analysis, MC38 tumor-bearing mice received a single intravenous injection of 1 × 10⁷ CFU per mouse of EcN-SAS or attenuated EcN^Δ^*^pgi^*-SAS. On days 1, 2, 4, 6 and 8 post-injection (n = 5 mice per time point per strain), mice were euthanized. Hearts, livers, spleens, lungs, kidneys, and tumors were harvested, weighed, homogenized in sterile PBS, and the resulting tissue homogenates were serially diluted and plated on LB agar containing appropriate antibiotics. Colonies were counted after overnight incubation at 37 °C, and the number of CFU per gram of tissue was calculated.

### Immune cell-mediated cytotoxicity against desialylated tumor cells

Human peripheral blood mononuclear cells (hPBMCs) were isolated from the whole blood of healthy donors by density gradient centrifugation using Ficoll-Paque PLUS (Cytiva). NK92 cells were obtained from ATCC and cultured in MEM-α supplemented with 12.5% FBS, 12.5% horse serum, 1% penicillin/streptomycin, and 100 U ml⁻¹ recombinant IL-2. Tumor cells were pre-incubated with different engineered bacteria for 4 h, washed extensively with PBS to remove unbound bacteria, and co-cultured with hPBMCs or NK92 cells at effector-to-target (E:T) ratio of 5:1 for an additional 4 h. Cytotoxicity of immune cells against tumor cells was assessed by measuring lactate dehydrogenase (LDH) release using CytoTox 96 Non-Radioactive Cytotoxicity Assay Kit (Promega) according to the manufacturer’s protocol. Absorbance was measured at 490 nm using a BioTek microplate reader.

### Macrophage-mediated phagocytosis of tumor cells

SK-BR-3 cells were pre-treated with EcN^Δpgi^, EcN^Δpgi^-SS, EcN^Δpgi^-S^E647A^S, or EcN^Δpgi^-SAS at a 1:10 ratio for 4 h as described above. After washing, tumor cells were labeled with 5 μM carboxyfluorescein succinimidyl ester (CFSE) for 15 min at 37 °C, washed, and co-cultured with human THP-1-derived macrophages, primary mouse bone marrow-derived macrophages (BMDMs), or RAW 264.7 cells at a macrophage: tumor ratio of 5:1 for 8 h. Macrophages were stained with APC-conjugated anti-CD11b antibody (BioLegend) prior to analysis. Phagocytosis was assessed by immunofluorescence microscopy and quantified by flow cytometry (Beckman Coulter), with phagocytosis defined as CFSE-positive, CD11b-APC-positive double-positive events. For macrophage desialylation and functional assay, THP-1-derived macrophages were pre-treated with EcN^Δpgi^-SS or EcN^Δpgi^-S^E647A^S at a macrophage-to-bacteria ratio of 1:10 for 4 h, followed by extensive PBS washing. The pre-treated macrophages were then co-cultured with CFSE-labeled SK-BR-3 tumor cells at an E: T ratio of 5:1 for 8 h. Tumor cell phagocytosis was subsequently assessed by flow cytometry and further visualized by confocal microscopy.

### Synergistic inhibition of subcutaneous tumor growth with PD-L1 antibody

For subcutaneous tumor xenograft experiments, 2 × 106 4T1-HER2 cells were injected subcutaneously into the right flank of BALB/c mice. Tumor volumes were measured every 2-3 days with digital calipers and calculated as (length × width²)/2. When tumors reached a volume of 50-100 mm³, mice were randomized. To assess synergistic therapeutic efficacy, mice bearing established tumors were divided into treatment groups (n = 5 per group). Bacteria (1 × 10^5^ CFU in 100 μl of PBS per mouse) were administered intravenously via the tail vein every three days for a total of five doses. Anti-PD-L1 antibody (clone 10F.9G2, BioXCell) was administered intraperitoneally at 50 μg per mouse in 100 μl PBS every three days for a total of five doses. Mice were monitored twice a day for body weight and tumor volume. Mice were euthanized on day 21 post treatment. Cytokine (IFNγ, TNFα, IL-12p70) were measured by commercial ELISA kits (Sigma-Aldrich), all according to the manufacturer’s protocol. For imaging of desialylation in tumor tissues, frozen tissues were embedded in optimal cutting temperature (OCT) compound, cut into 8-μm thick sections, fixed in 4% paraformaldehyde, blocked with 5% BSA in PBS, and stained with SNA-Cy5 (1:200 dilution) and DAPI (1 μg mL⁻¹). The sfGFP reporter expressed in engineered bacteria allows visualization of bacterial distribution in tissue sections. Images were acquired on a Zeiss LSM 880 confocal microscope. Fluorescence intensity was quantified using ImageJ in regions of interest (ROIs) defined by bacterial localization.

### Flow cytometry of tumor-infiltrating immune cells

The subcutaneous tumors were harvested, minced into small pieces, and digested with 1 mg mL⁻¹ collagenase IV (Sigma-Aldrich) and 20 U mL⁻¹ DNase I (Roche) in RPMI-1640 for 30 min at 37°C with continuous shaking. Single-cell suspensions were passed through 70-μm cell strainers, washed with PBS, and incubated with red blood cell lysis buffer (BioLegend) for 5 min on ice. Cells were blocked with anti-CD16/32 antibody (1:100, BioLegend) for 10 min at 4°C, then stained with cocktails of fluorochrome-conjugated antibodies for 30 min at 4°C. For intratumoral staining of lymphocytes, the antibody panel was as followed: CD45-Alexa Fluor®700, CD3-APC, CD4-FITC, CD8-PE/Cy7, and NKp46-PE (all from BioLegend). For intratumoral staining of myeloid cells, the antibody panel was as followed: CD45-Alexa Fluor®700, CD80-Brilliant Violet 510, CD206-APC, F4/80-PE/Cy7, Ly-6C-Brilliant Violet 421 (all from BioLegend). For intratumoral Treg cells staining, surface markers were stained first, followed by fixation and permeabilization before intracellular Foxp3 staining. The antibody panel was as follows: CD45-Alexa Fluor®700, CD4-PE (surface), and Foxp3-APC (intracellular, all from BioLegend). Cells were fixed and permeabilized using the Foxp3/Transcription Factor Staining Buffer Set (eBioscience) according to the manufacturer’s instructions. 7AAD viability staining (BioLegend) was used to exclude dead cells. Samples were acquired on a CytoFLEX flow cytometer (Beckman Coulter) and analyzed with FlowJo software (v10.8.1). Gating strategies are provided in Supplementary Fig.**27-30.**

### Quantification of cell surface sialylation levels on tumor cells

In vivo desialylation of tumor cell surface was assessed in established 4T1-HER2 subcutaneous tumors. Subcutaneous tumors were harvested, and single-cell suspensions were prepared with red blood cells lysis performed as described above. Cells were stained with fluorophore-conjugated anti-CD45 antibody to distinguish immune cells, and CD45⁻ tumor cells were gated for downstream analysis. Tumor cell surface sialic acids were stained with SNA-Cy5 (1:200 dilutions, 30 min at 4 °C) and quantified by flow cytometry on a CytoFLEX instrument. The mean fluorescence intensity (MFI) was normalized to that of PBS-treated controls.

### Quantification of liberated sialic acids in the tumor microenvironment

Tumors were harvested, minced into small pieces, and digested into single-cell suspensions as described above. The resulting single-cell suspension was centrifuged at 500 × g for 10 min at 4 °C, and the supernatant was collected from each sample. Proteins were precipitated by adding an equal volume of ice-cold methanol, followed by ultracentrifugation to remove the precipitated protein aggregates. The clarified supernatant was then filtered through a 0.22 μm membrane and subjected to LC-MS analysis in selected ion monitoring (SIM) mode.

### Inhibition of colorectal cancer in AOM/DSS-treated mice

Female C57BL/6 mice (6-8 weeks old) were injected intraperitoneally with 10 mg kg⁻¹ azoxymethane (AOM). After one week, colitis was induced by supplementing 3% (w/v) dextran sulfate sodium (DSS) to autoclaved drinking water for 7 days, followed by 7 days of normal autoclaved drinking water. This cycle was repeated five times to establish a colorectal cancer (CRC) mouse model. Healthy control mice received AOM alone. The AOM/DSS-induced CRC-bearing mice were randomized into treatment groups (n = 6 per group) and treated orally with 1 × 10⁹ CFU per mouse of the indicated bacterial strains, or an equal volume of PBS every three days for a total of five doses. At the experimental endpoint, colons were excised, measured for length, and processed for histology (H&E staining), immunofluorescence analysis, and bacterial enumeration.

### Histology and immunofluorescence of colon tissues

The collected mouse colon tissues were fixed in 4% paraformaldehyde, embedded in paraffin, sectioned into 5-μm thick sections, and stained with haematoxylin and eosin (H&E). Histological scores were assessed in a blinded manner based on crypt damage, inflammation, and tumor infiltration. For immunofluorescence, paraffin sections were deparaffinized, rehydrated, and subjected to heat-induced antigen retrieval in citrate buffer (pH 6.0). For imaging of desialylation in mouse colon sections, sections were stained with SNA-Cy5 (1:200 dilution). sfGFP expressed by engineered bacteria enables direct in situ visualization of bacterial distribution within colon tissue. For analysis of mouse intestinal structure and barrier integrity, colon sections were blocked with 5% donkey serum and stained overnight at 4 °C with primary antibodies: anti-ZO-1 (1:200 dilution), anti-Occludin (1:200 dilution), anti-Mucin 2 (1:200 dilution), anti-E-cadherin (1:200 dilution), and anti-Ki67 (1:200 dilution). After washing, sections were incubated with Alexa Fluor-conjugated secondary antibodies (1:500 dilution) for 1 h at room temperature, counterstained with DAPI, and imaged by confocal microscopy.

### Statistical analysis

Data are presented as the mean ± s.d. unless otherwise indicated. Statistical analyses were performed using GraphPad Prism 9.0. Comparisons between two groups were made using a two-tailed unpaired Student’s t-test. Comparisons between multiple groups were performed using one-way or two-way analysis of variance (ANOVA) with appropriate post-hoc multiple comparison tests. Survival curves were analyzed using the log-rank (Mantel-Cox) test. Normality of data distribution was assessed using the Shapiro–Wilk test. Homogeneity of variances was assessed using Levene’s test. P < 0.05 was considered statistically significant. Exact P values are provided in the figure legends where applicable.

## Data availability

All data supporting the results are available in the main text or the supplementary materials. Additional data are available from the corresponding author upon reasonable request. Source data are provided with this paper.

## Acknowledgements

This work was supported by the National Key R&D Program of China (2022YFA1104900); National Natural Science Foundation of China (22677209, 22293053, 22022706 and 92353301) and GBRCE for Functional Molecular Engineering.

## Author contributions

J.W., J.W., Z.-W.M. and W.X. conceived and designed the study. J.W., J.W., T.O., B.S and H.M. performed in vitro characterization. J.W., J.W. and B.S. performed in vivo experiments. J.W., J.W. and W.X. wrote the manuscript.

## Additional information

Supplementary information the online version contains supplementary material available at

