## Supplemental information for "Targeted tumor desialylation by engineered probiotic reprograms the tumor microenvironment to potentiate immunotherapy"

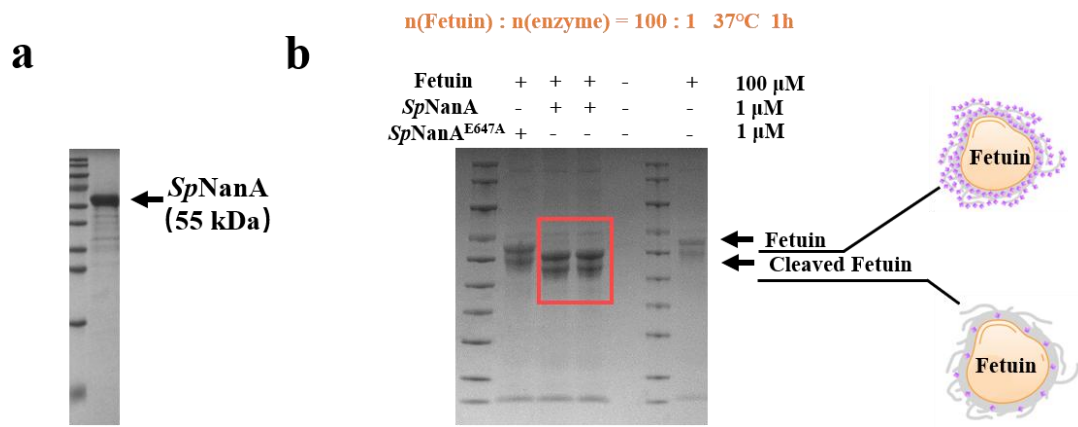

**Supplementary Fig. 1 SDS-PAGE analysis of purified *SpNanA*, and cleaved fetuin. a**, Purified recombinant *SpNanA* was resolved by SDS-PAGE and visualized via Coomassie blue staining. **b**, Fetuin was incubated with *SpNanA* or the catalytically inactive mutant *SpNanA*<sup>E647A</sup> at a 1:100 for 1 h at 37°C. Reaction products were analyzed by SDS-PAGE.

**n(Fetuin) : n(enzyme) = 100 : 1 37°C 1h**

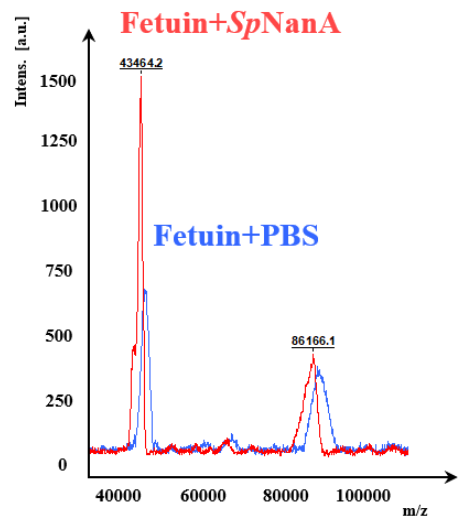

**Supplementary Fig. 2 MALDI-TOF mass spectrometry analysis of *SpNanA*-treated fetuin.** Fetuin was incubated with *SpNanA* (red trace) or left untreated (blue trace) at 37 °C for 1 h, and reaction products were analyzed using MALDI-TOF mass spectrometry. Left peaks correspond to monomeric fetuin, and right peaks correspond to dimeric fetuin. Data are representative of three independent experiments.

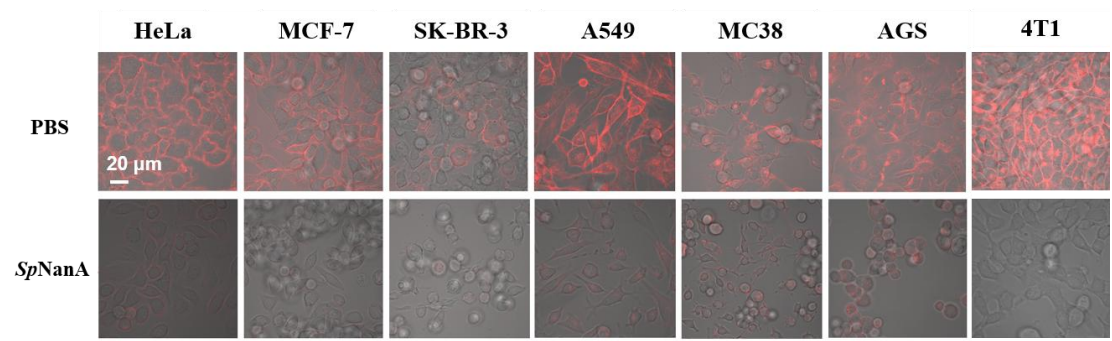

**Supplementary Fig. 3** The indicated cell lines (HeLa, MCF7, SK-BR-3, A549, MC38, AGS and 4T1) were incubated with 10 μM purified *SpNanA* at 37 °C for 0 min (top row) or 30 min (bottom row). Cells were stained with SNA-Cy5 to label cell-surface sialic acids. Scale bar, 20 μm.

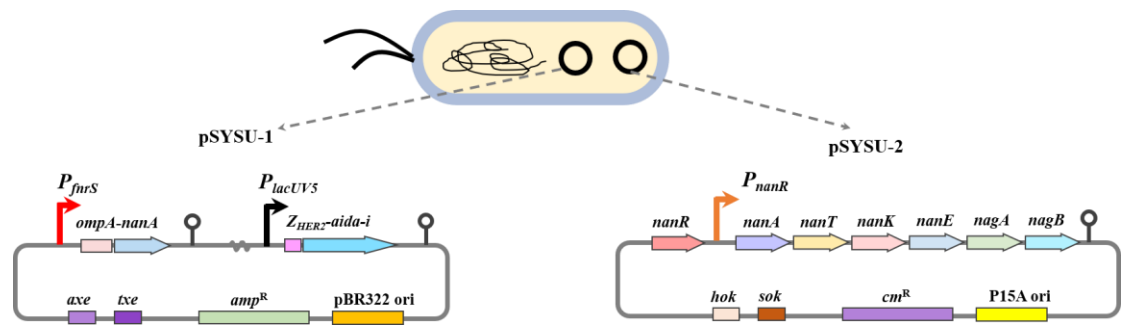

**Supplementary Fig. 4** Plasmid pSYSU-1 contains the following elements: a hypoxia-inducible $P_{fnrS}$ promoter driving transcription of the gene encoding the OmpA-NanA fusion protein (*ompA-nanA*), enabling surface display of *SpNanA*; a constitutive $P_{lacUV5}$ promoter driving transcription of the gene encoding the anti-HER2 affibody-AIDA-I fusion protein (*Z<sub>HER2</sub>-aida-i*), enabling its surface display; the Axe/Txe plasmid stability system (*axe/txe*); the pBR322 origin of replication (pBR322 ori); and an ampicillin resistance gene ($amp^R$). Plasmid pSYSU-2 contains: a constitutively expressed NanR repressor (*nanR*); a sialic acid-inducible $P_{nanR}$ promoter driving transcription of the catabolic gene cluster (*nanT*, *nanA*, *nanK*, *nanE*, *nagA* and *nagB*); the Hok/Sok (*hok/sok*) plasmid stability system; the P15A origin of replication; and chloramphenicol resistance gene ($cm^R$).

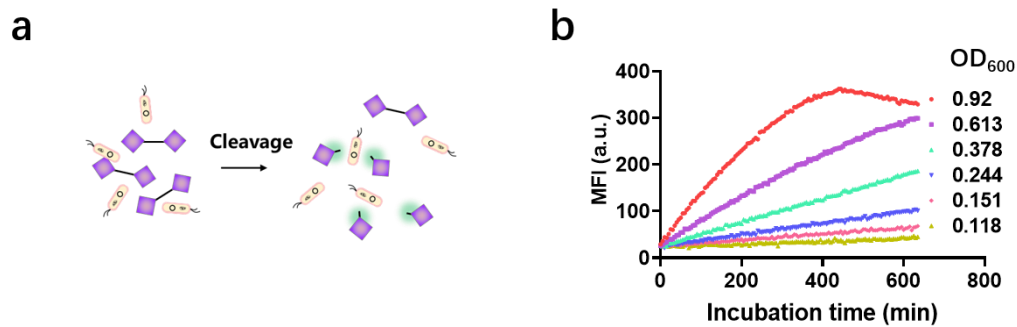

**Supplementary Fig. 5 Qualitative assessment of sialidase activity in engineered bacteria using a commercial sialidase assay kit.** **a**, Schematic depicting cleavage of the fluorogenic substrate by engineered bacteria, resulting in fluorescence emission. **b**, Time-dependent increase in fluorescence upon cleavage of excess substrate by engineered bacteria at varying cell densities (quantified by OD<sub>600</sub>). Fluorescence intensity was measured at the indicated time points. Data are representative of three independent experiments.

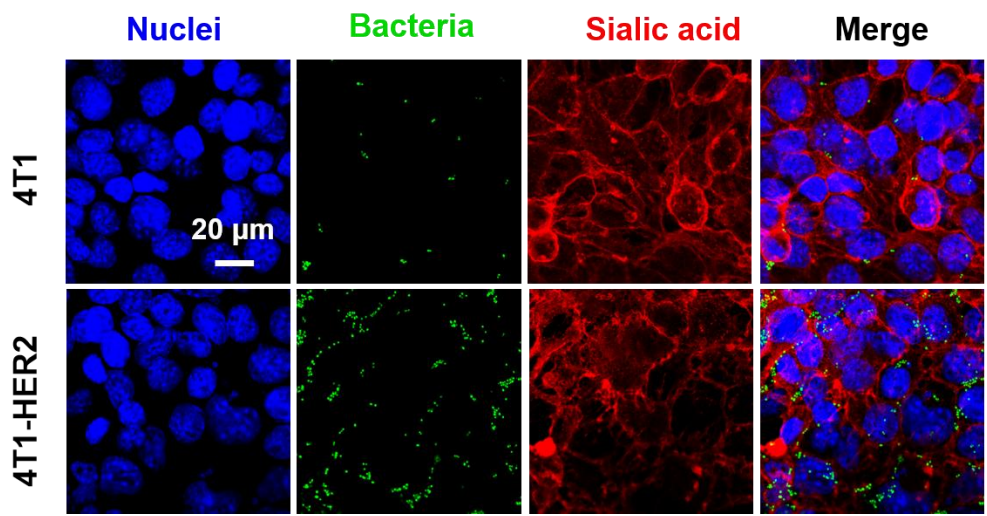

**Supplementary Fig. 6 Confocal imaging of bacteria targeting to 4T1 and 4T1-HER2 cells.** Engineered bacteria expressing sfGFP were co-cultured with tumor cells at a 100:1 bacteria-to-cell ratio for 4 h at 37 °C. Following washing, cells were stained with Hoechst for nuclei and SNA-Cy5 for cell-surface sialic acids. Blue: nuclei; green: bacteria; red: sialic acid. Scale bar, 20 μm.

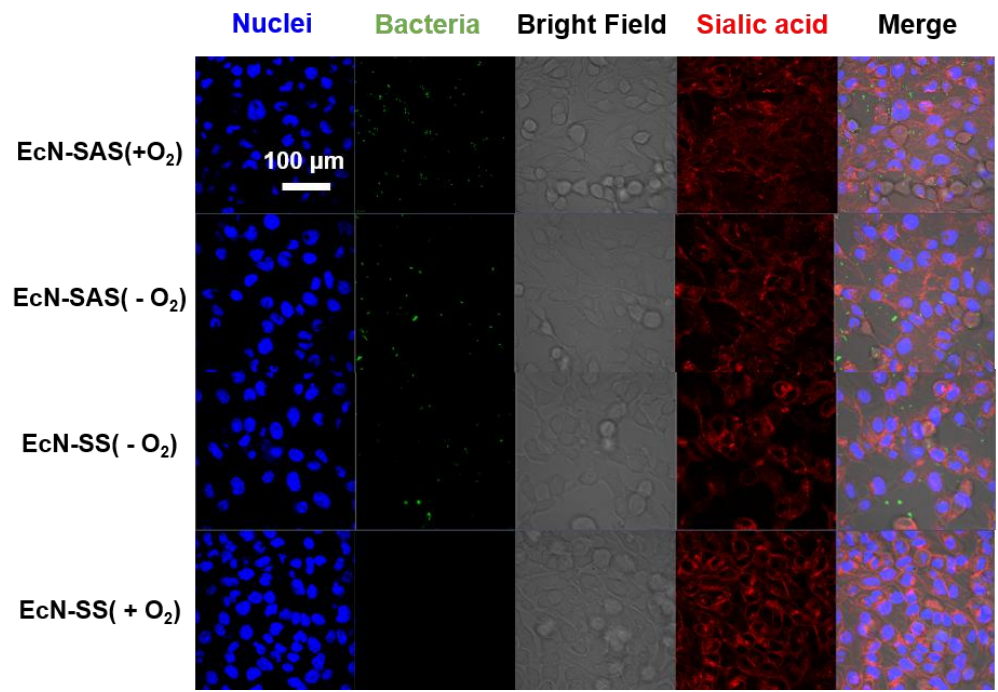

**Supplementary Fig. 7 Confocal imaging of bacteria targeting to 4T1 cells under normoxic and hypoxic conditions.** Engineered bacteria (EcN-SAS and EcN-SS) expressing sfGFP were co-cultured with 4T1 cells at a 100:1 bacteria-to-cell ratio for 4 h at 37 °C under normoxic or hypoxic conditions. Following washing, cells were stained with Hoechst for nuclei and SNA-Cy5 for cell-surface sialic acids. Blue: nuclei; green: bacteria; red: sialic acid. Scale bar, 100 μm.

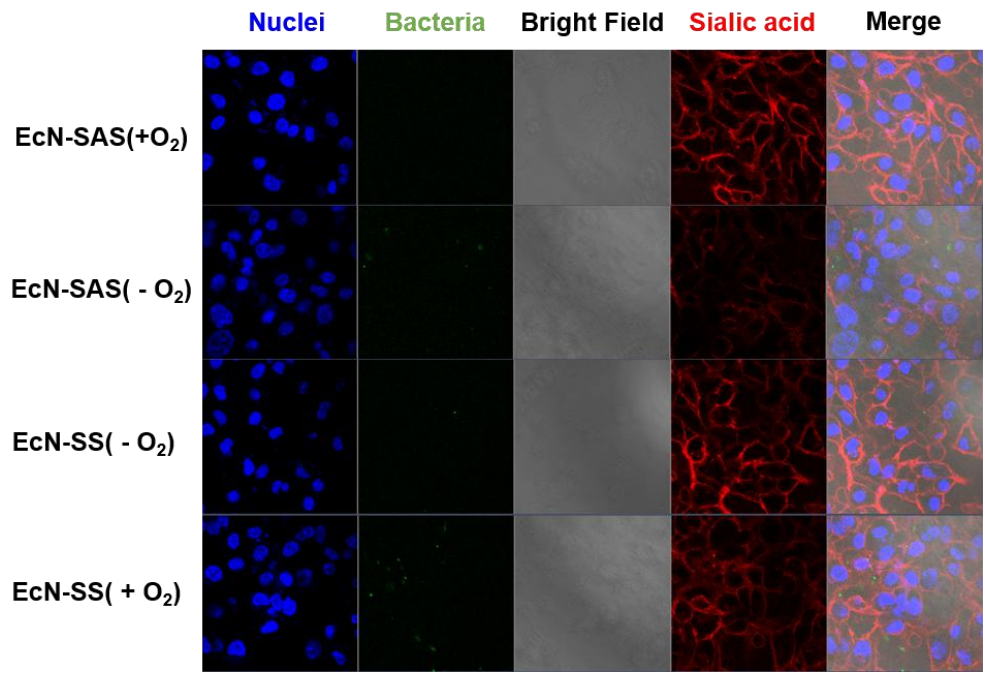

**Supplementary Fig. 8 Confocal imaging of bacteria targeting to MDA-MB-231 cells under normoxic and hypoxic conditions.** Engineered bacteria (EcN-SAS and EcN-SS) expressing sfGFP were co-cultured with MDA-MB-231 cells at a 100:1 bacteria-to-cell ratio for 4 h at 37 °C under normoxic or hypoxic conditions. Following washing, cells were stained with Hoechst for nuclei and SNA-Cy5 for cell-surface sialic acids. Blue: nuclei; green: bacteria; red: sialic acid. Scale bar, 100 μm.

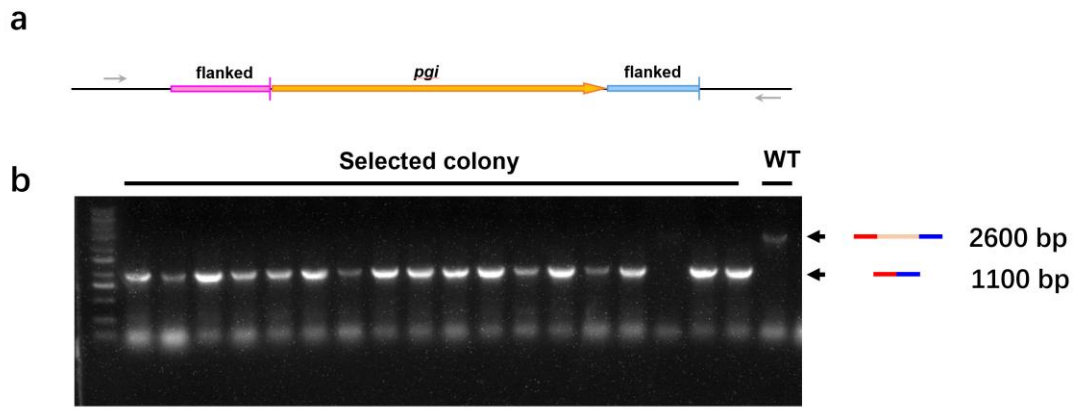

**Supplementary Fig. 9 Genomic deletion of *pgi* in EcN.** **a**, Schematic showing Sanger sequencing validation of *pgi* deletion in the EcN genome. **b**, Colony PCR validation of *pgi* deletion. The forward primer was designed upstream of the 5' homology arm, and the reverse primer was designed downstream of the 3' homology arm. Successful deletion generated a 1,107 bp PCR amplicon, while the wild-type locus yielded a 2,761 bp product.

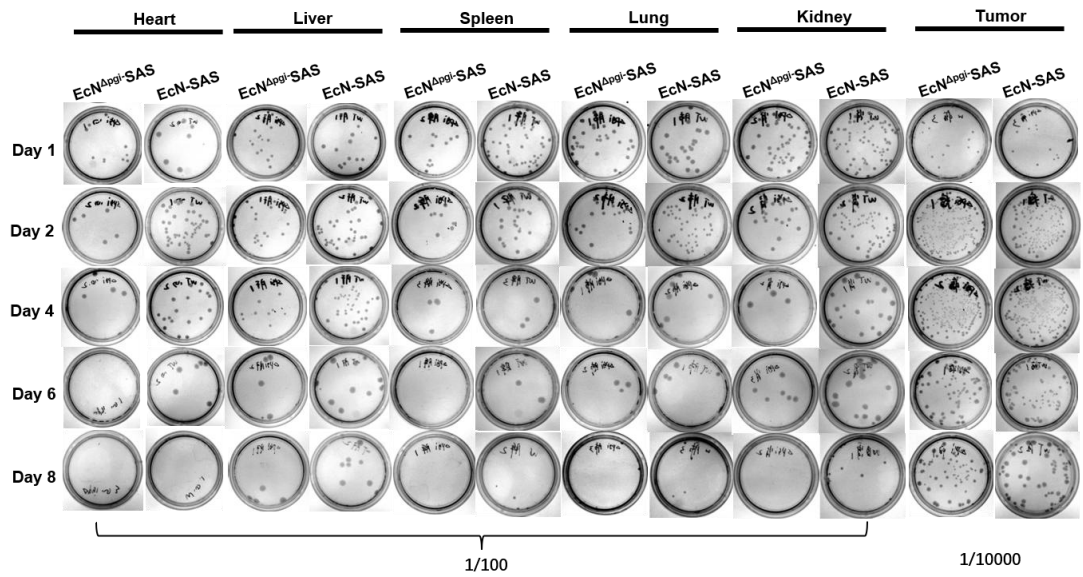

**Supplementary Fig. 10 Bacterial colonization in mouse organs and tumors.** Tissue homogenates collected from the heart, liver, spleen, lung, kidney and tumor were plated onto LB agar for colony enumeration. Homogenates of the heart, liver, spleen, lung and kidney were diluted 100-fold before plating, while tumor homogenates were diluted 10,000-fold. Data are representative of three independent experiments.

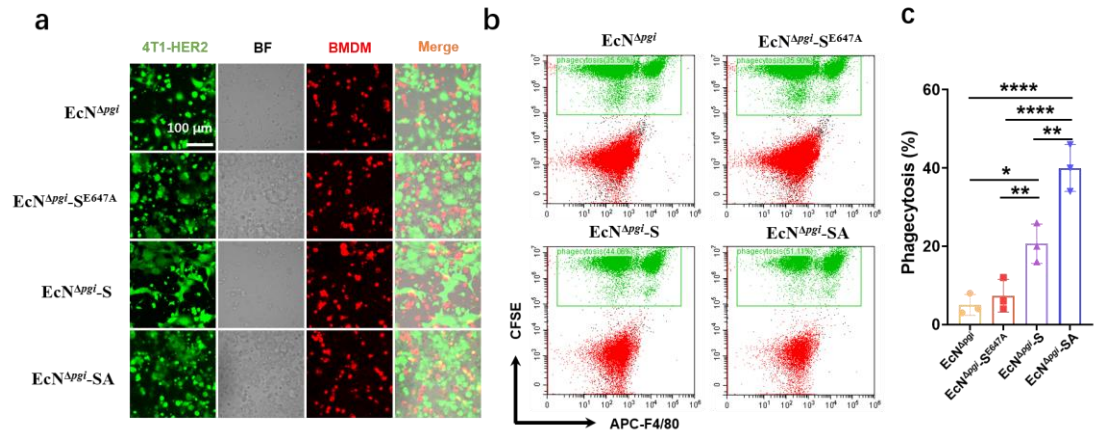

**Supplementary Fig. 11 Phagocytosis of tumor cells by BMDM macrophages after co-culture with engineered bacteria.** **a**, Confocal images showing phagocytosis of CFSE-labelled tumor cells (green) by BMDM macrophages stained with APC-conjugated anti-F4/80 antibody (red). Tumor cells were pre-incubated with the indicated bacterial strains prior to co-culture with BMDM macrophages. **b**, Representative flow cytometry histograms gated on F4/80<sup>+</sup> macrophages showing CFSE fluorescence intensity, which reflects the extent of tumor cell phagocytosis across bacterial treatment groups. **c**, Quantification of phagocytosis based on flow cytometry data. Data are presented as mean ± s.d. from $n = 3$ biological replicates. Statistical analysis was performed using one-way ANOVA with Tukey's multiple comparisons test. \*$P < 0.05$, \*\*$P < 0.01$, \*\*\*$P < 0.001$ and \*\*\*\*$P < 0.0001$.

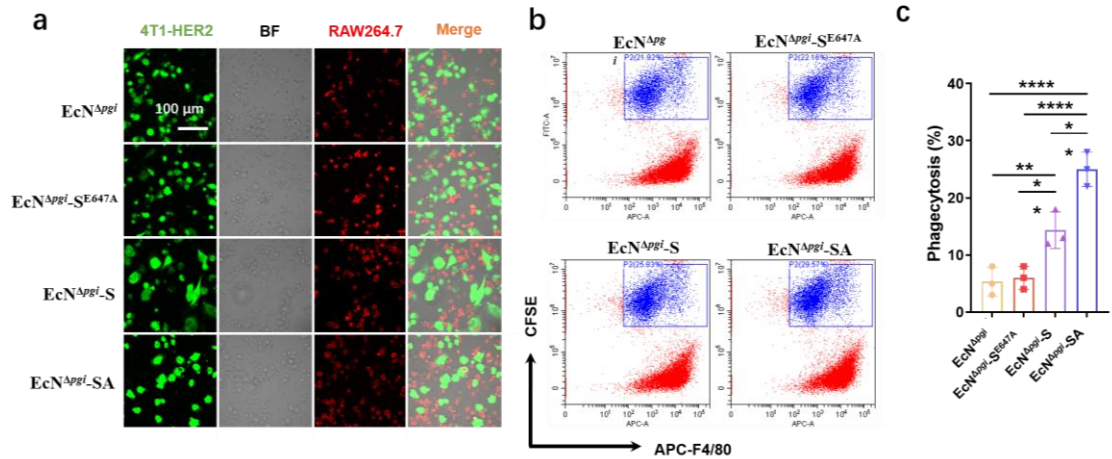

**Supplementary Fig. 12 Phagocytosis of tumor cells by RAW 264.7 macrophages after co-culture with engineered bacteria.** **a**, Confocal images showing phagocytosis of CFSE-labelled tumor cells (green) by RAW 264.7 macrophages stained with APC-conjugated anti-F4/80 antibody (red). Tumor cells were pre-incubated with the indicated bacterial strains prior to co-culture with RAW 264.7 macrophages. **b**, Representative flow cytometry histograms gated on F4/80<sup>+</sup> macrophages showing CFSE fluorescence intensity, which reflects the extent of tumor cell phagocytosis across bacterial treatment groups. **c**, Quantification of phagocytosis based on flow cytometry data. Data are presented as mean ± s.d. from $n = 3$ biological replicates. Statistical analysis was performed using one-way ANOVA with Tukey's multiple comparisons test. \*$P < 0.05$, \*\*$P < 0.01$ and \*\*\*\*$P < 0.0001$.

**PDL1 (*i.p.*) injection
on Day 9, 12, 15, 18**

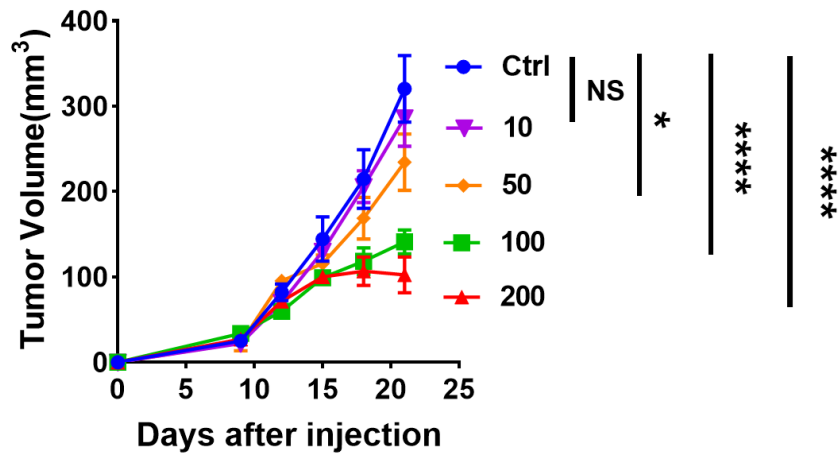

**Supplementary Fig. 13 Dose titration of anti-PD-L1 antibody in the subcutaneous 4T1-HER2 tumor model.** Mice bearing subcutaneous 4T1-HER2 tumors received anti-PD-L1 antibody at doses of 10, 50, 100 or 200 µg per mouse via intraperitoneal injection, administered every three days for a total of four doses. Tumor volumes were monitored to evaluate therapeutic efficacy and select a suboptimal dose for subsequent combination studies. Data are presented as mean ± s.d. from $n = 5$ biologically independent mice per group. Statistical analysis was performed using one-way ANOVA followed by Tukey's multiple comparisons test. \*$P < 0.05$ and \*\*\*\*$P < 0.0001$. NS, not significant.

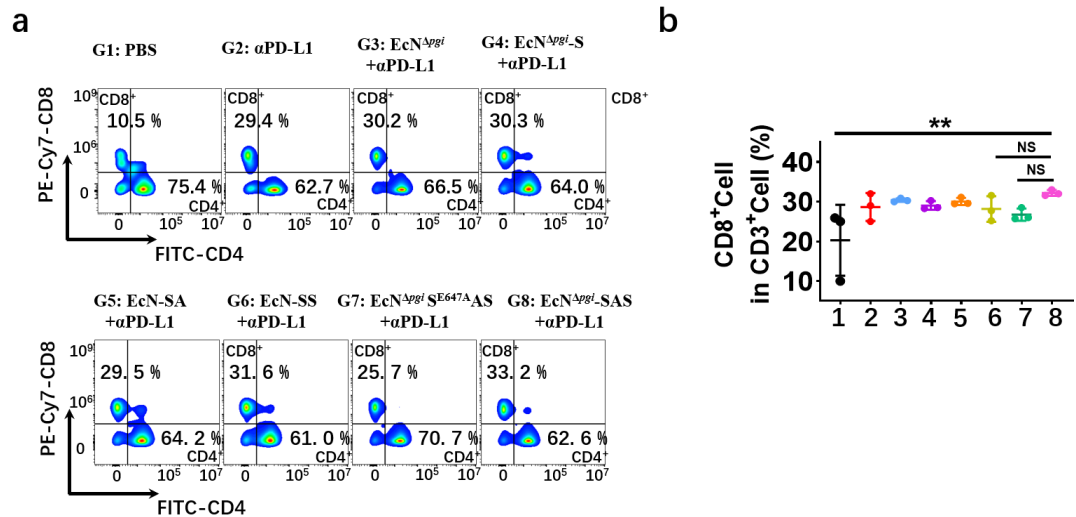

**Supplementary Fig. 14 Flow cytometric analysis of CD8<sup>+</sup> T-cell infiltration in tumor tissues. a,** Representative flow cytometry plots showing CD4 and CD8 expression, gated in CD3<sup>+</sup> cells isolated from tumors of mice across groups G1-G8. CD4 was detected on the FITC channel, and CD8 in the PE-Cy7 channel. **b,** Quantification of CD8<sup>+</sup> T-cell frequency within CD3<sup>+</sup> tumor-infiltrating lymphocytes across treatment group. Data are presented as mean ± s.d. from $n = 3$ biologically independent mice per group. Statistical analysis was performed using one-way ANOVA with Tukey's multiple comparisons test. $**P < 0.01$. NS, not significant.

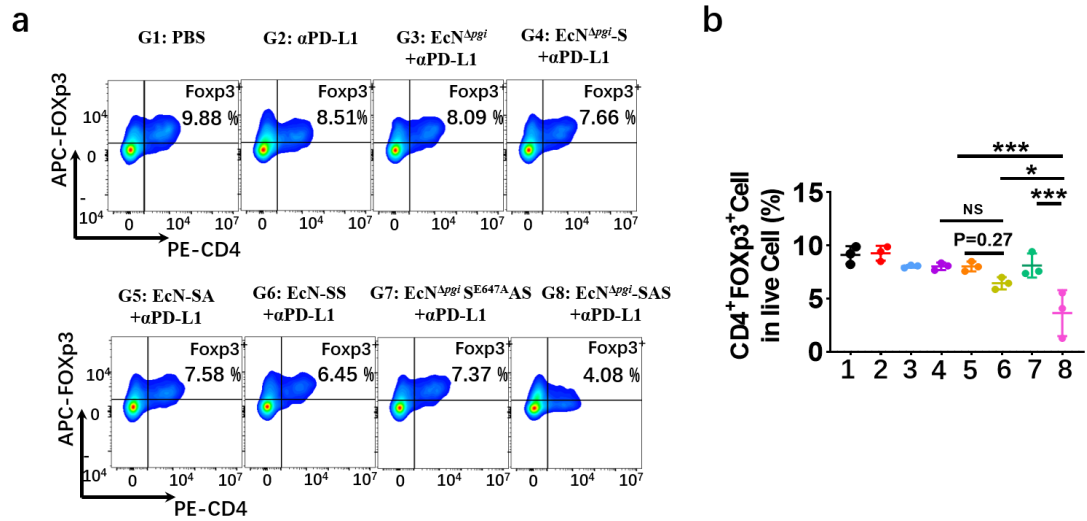

**Supplementary Fig. 15 Flow cytometric analysis of CD4<sup>+</sup>FOXP3<sup>+</sup> Treg cell infiltration in tumor tissues.** **a**, Representative flow cytometry plots showing CD4 and FOXp3 expression, gated on CD45<sup>+</sup> cells isolated from tumors of mice across groups G1-G8. CD4 was detected in the PE channel, and FOXP3 in the APC channel. **b**, Quantification of CD4<sup>+</sup>FOXP3<sup>+</sup> T cell frequency within CD45<sup>+</sup> cells across treatment group. Data are presented as mean ± s.d. from n = 3 mice per group. Statistical analysis was performed using one-way ANOVA with Tukey's multiple comparisons test. \**P* < 0.05 and \*\*\**P* < 0.001. NS, not significant.

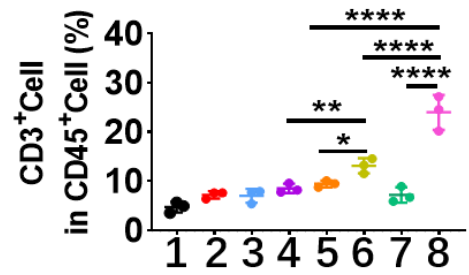

**Supplementary Fig. 16 Flow cytometric analysis of CD3<sup>+</sup> T-cell infiltration in tumor tissues.** Quantification of CD3<sup>+</sup> T-cell frequency within CD45<sup>+</sup> tumor-infiltrating lymphocytes across treatment group. Data are presented as mean ± s.d. from $n = 3$ biologically independent mice per group. Statistical analysis was performed using one-way ANOVA with Tukey's multiple comparisons test. \*$P < 0.05$, \*\*$P < 0.01$ and \*\*\*\*$P < 0.0001$. NS, not significant.

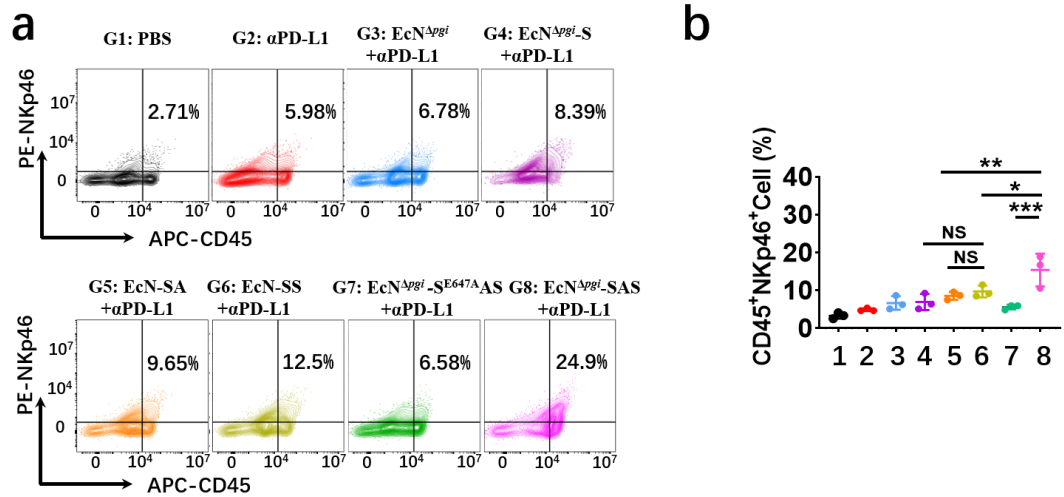

**Supplementary Fig. 17 Flow cytometric analysis of NK cell infiltration in tumor tissues. a,** Representative flow cytometry plots showing CD45<sup>+</sup>NKp46<sup>+</sup> expression gated on live cells isolated from tumor tissues of mice in groups G1-G8. CD45 was detected in the APC channel, and NKp46 in the PE channel. **b,** Quantification of NK cell frequency among live cells for each treatment group. Data are presented as mean ± s.d. from n = 3 mice per group. Statistical analysis was performed using one-way ANOVA with Tukey's multiple comparisons test. \*$P$ < 0.05, \*\*$P$ < 0.01 and \*\*\*$P$ < 0.001. NS, not significant.

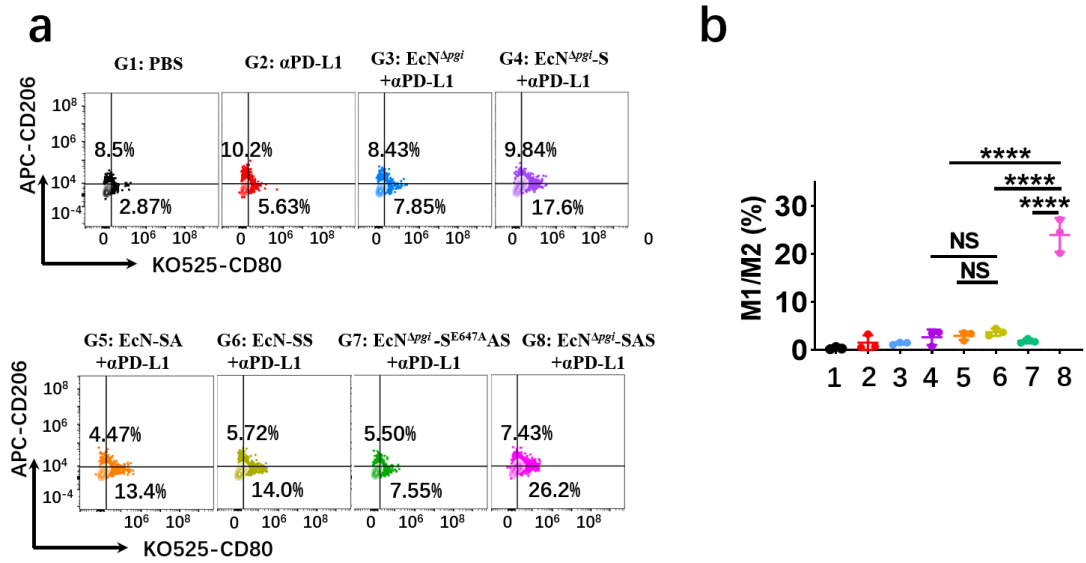

**Supplementary Fig. 18 Flow cytometric analysis of the M1/M2 macrophage ratio in tumor tissues.** **a**, Representative flow cytometry plots showing CD206 and CD80 expression gated on CD45<sup>+</sup>F4/80<sup>+</sup> cells isolated from tumor tissues of mice in groups G1-G8. CD80 was detected in the KO525 channel, and CD206 in the APC channel. **b**, Quantification of the M1/M2 macrophage ratio for each treatment group. Data are presented as mean ± s.d. from $n = 3$ mice per group. Statistical analysis was performed using one-way ANOVA with Tukey's multiple comparisons test. \*\*\*\*$P < 0.0001$; NS, not significant.

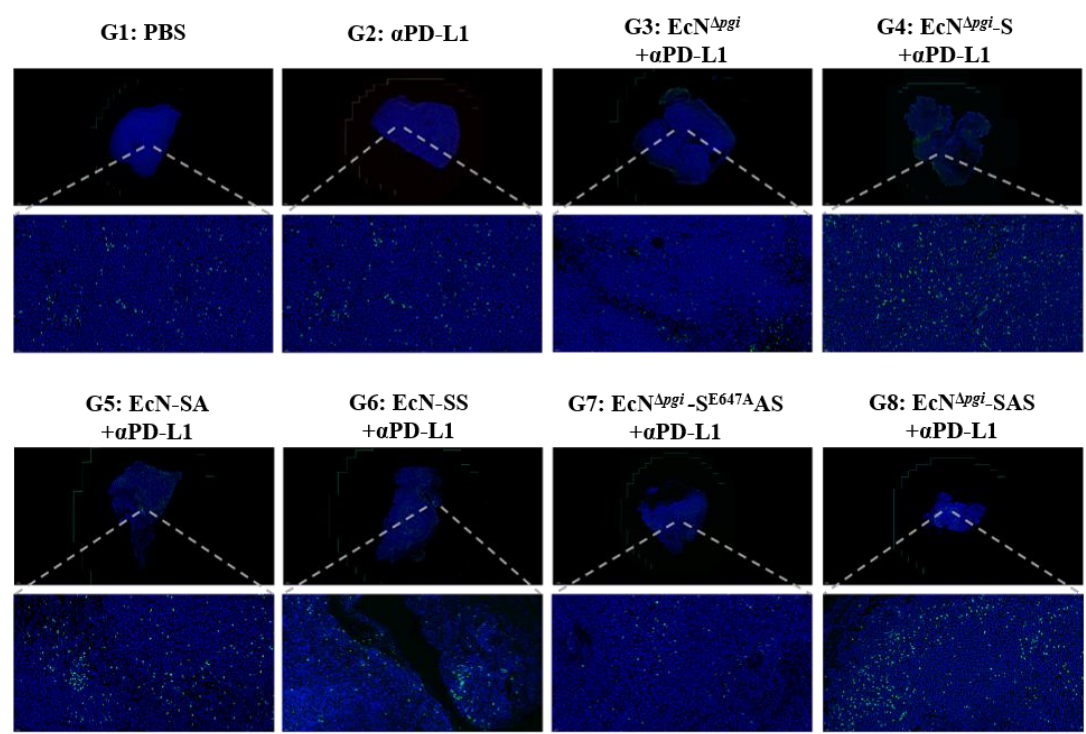

**Supplementary Fig. 19** Immunofluorescence staining assessing CD8<sup>+</sup> T cell infiltration in mouse tumor sections following the indicated treatments. Blue, nuclei; green, CD8<sup>+</sup> T cells.

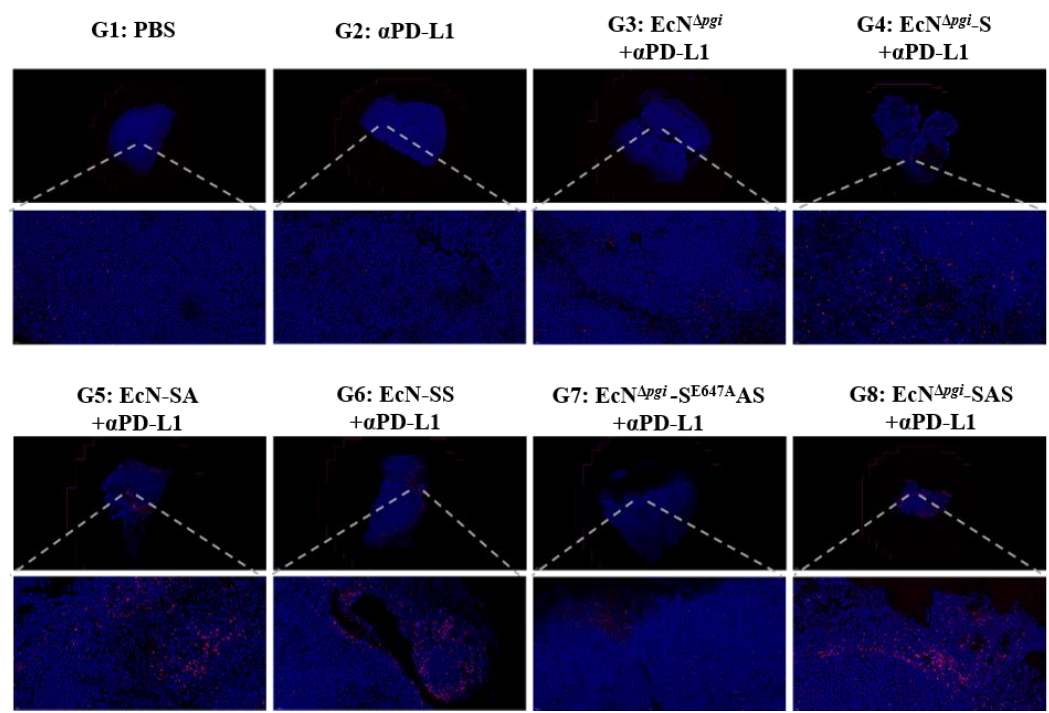

**Supplementary Fig. 20** Immunofluorescence staining assessing NK cell infiltration in mouse tumor sections following the indicated treatments. Blue, nuclei; red, NK cells.

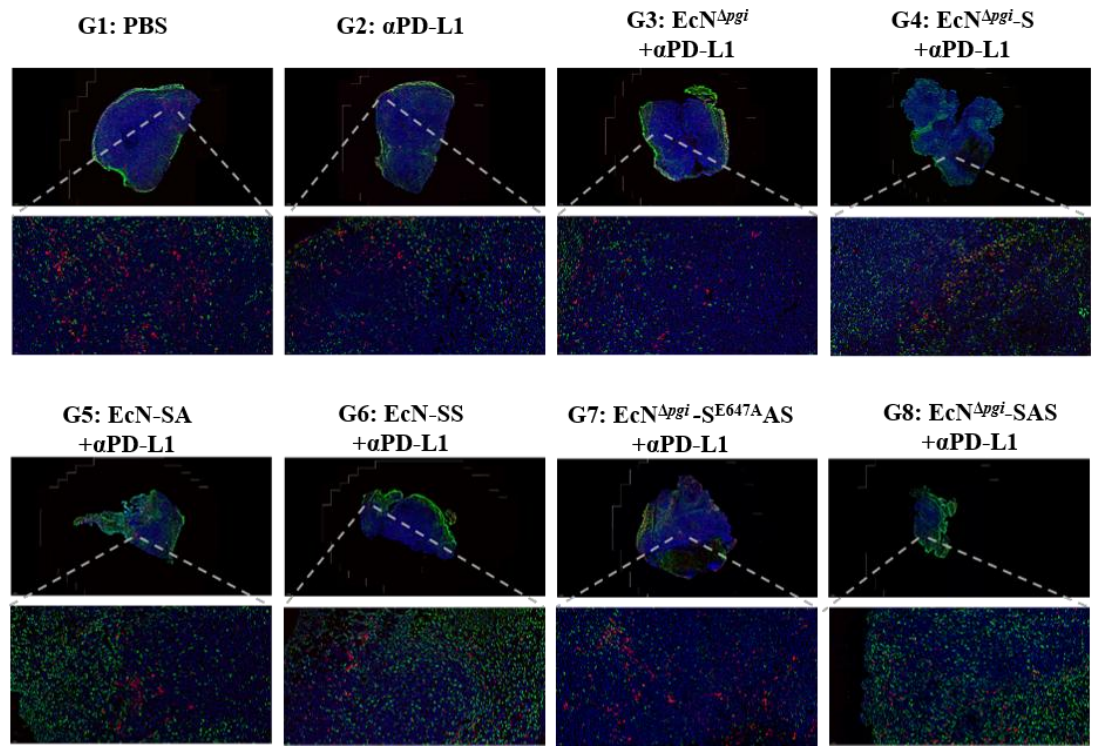

**Supplementary Fig. 21** Immunofluorescence staining assessing M1 and M2 macrophage infiltration in mouse tumor sections following the indicated treatments. Blue, nuclei; green, M1 macrophage; red: M2 macrophage.

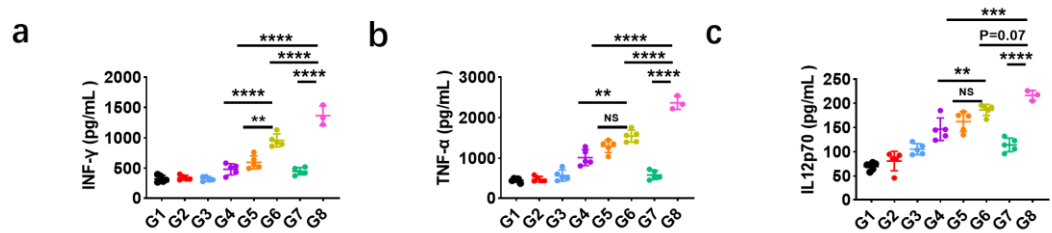

**Supplementary Fig. 22 Cytokine concentrations in mouse serum.** Concentrations of IFN-γ (a), TNF-α (b), and IL-12p70 (c) in serum collected from mice across the indicated treatment groups were quantified by ELISA. Data are presented as mean ± s.d. from $n = 5$ mice per group. Statistical analysis was performed using one-way ANOVA with Tukey's multiple comparisons test. \*$P < 0.05$, \*\*$P < 0.01$, \*\*\*$P < 0.001$, \*\*\*\*$P < 0.0001$; NS, not significant.

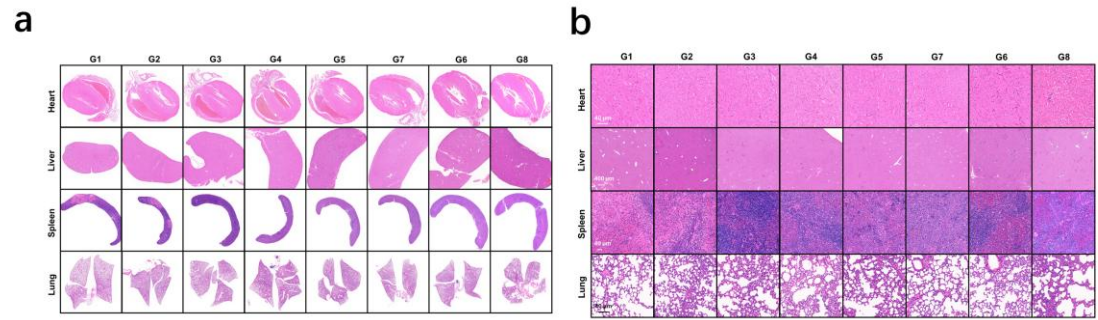

**Supplementary Fig. 23 H&E staining of heart, liver, spleen, lung, and kidney tissue sections from mice in groups G1–G8. a, Low-magnification overviews of tissue morphology. b, High-magnification micrographs of each organ. Scale bar, 40 μm.**

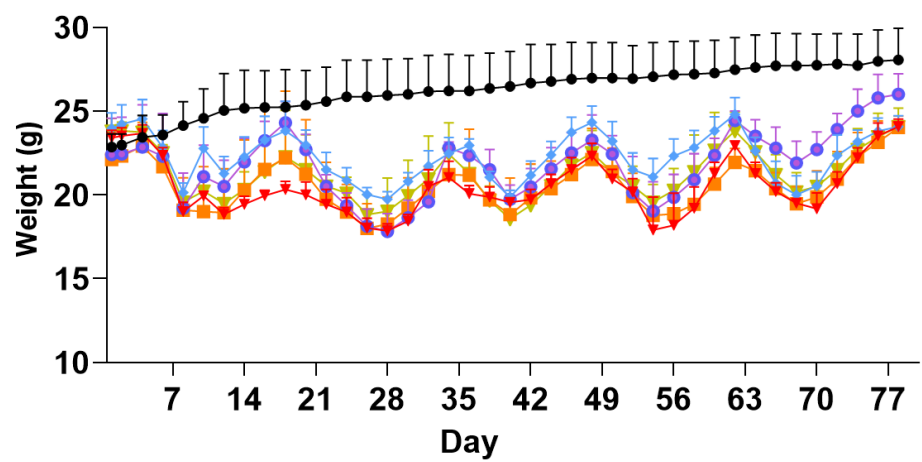

**Supplementary Fig. 24** Body weight changes in mice during colorectal cancer (CRC) induction. Body weights were monitored every two days throughout the induction period. Data are presented as mean ± s.d. from $n = 5$ mice per group.

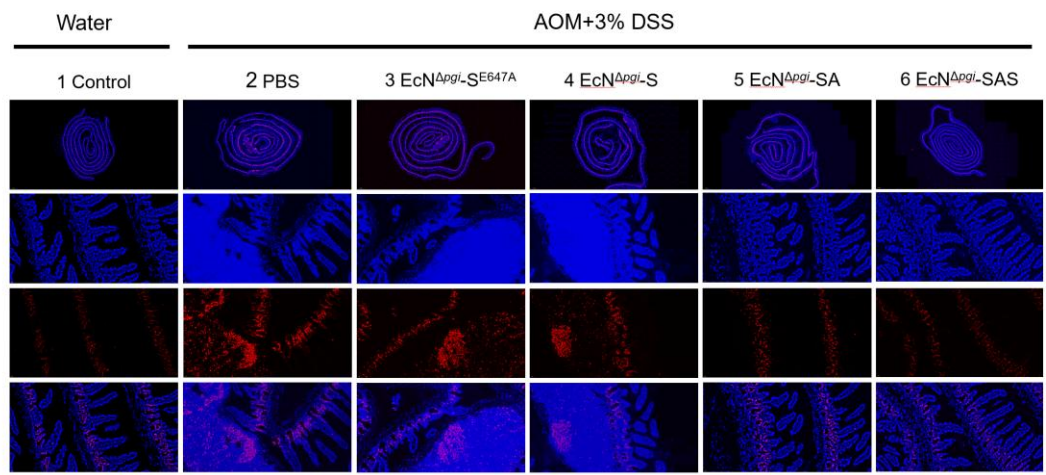

**Supplementary Fig. 25** Representative immunofluorescence staining images Ki67 expression in mice colon tissue sections after the indicated treatment. Blue: nuclei; red: Ki67.

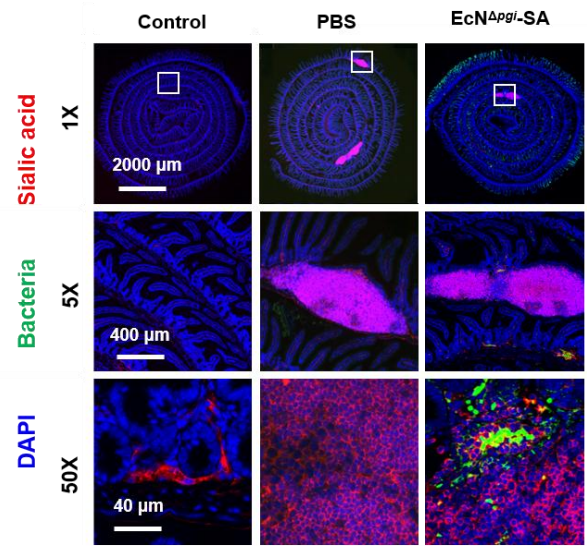

**Supplementary Fig. 26** Representative immunofluorescence staining images of bacteria and tumor sialylation in mice colon tissue sections after the indicated treatment. Red: sialic acid; green: bacteria; blue: nuclei.

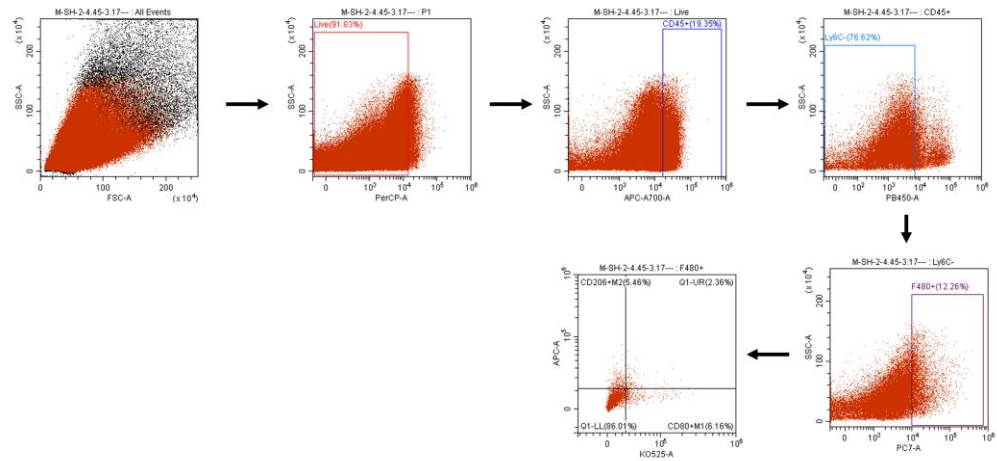

**Supplementary Fig. 27** Gating strategy for macrophage phenotype analysis by flow cytometry using mouse tumor tissues. Sequential gating steps were applied to identify live CD45<sup>+</sup>F4/80<sup>+</sup> macrophages, followed by sub-gating for CD80 (M1) and CD206 (M2) markers.

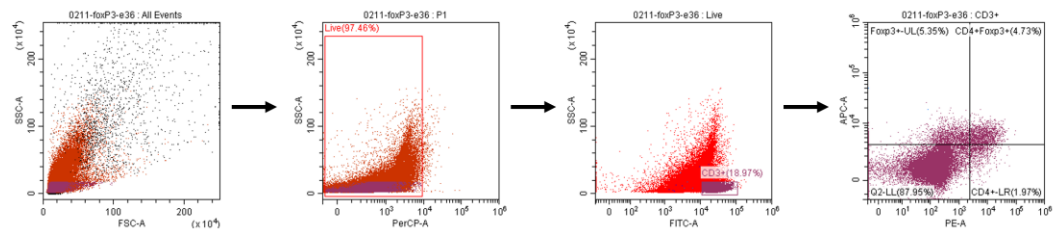

**Supplementary Fig. 28** Gating strategy for Foxp3<sup>+</sup> regulatory T cell analysis by flow cytometry using mouse tumor tissues. Sequential gating steps were used to identify live CD3<sup>+</sup>CD4<sup>+</sup>Foxp3<sup>+</sup> regulatory T cells.

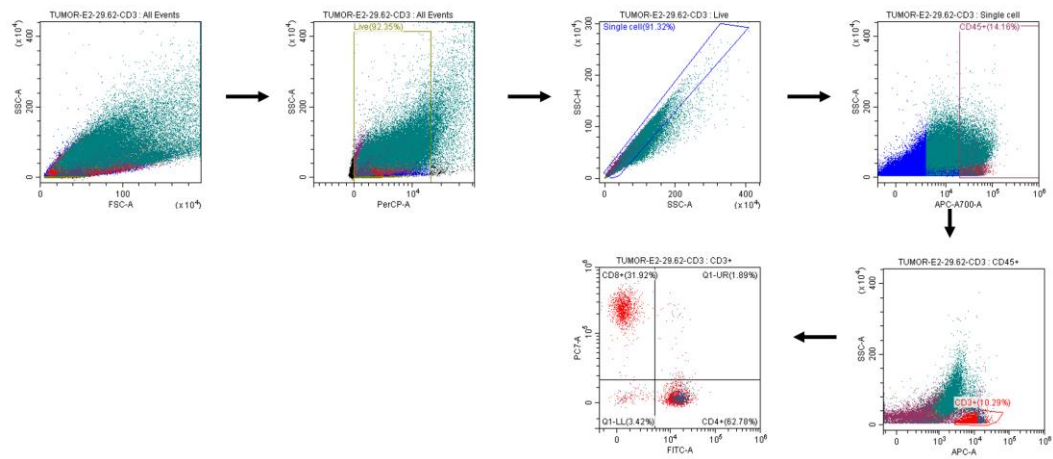

**Supplementary Fig. 29** Gating strategy for CD3<sup>+</sup> T cell analysis by flow cytometry using mouse tumor tissues. Sequential gating steps were applied to identify live CD45<sup>+</sup>CD3<sup>+</sup> T cells from tumor single-cell suspensions.

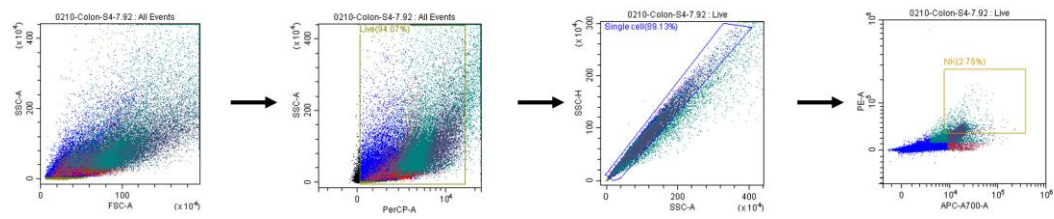

**Supplementary Fig. 30** Gating strategy for NK cell analysis by flow cytometry using mouse tumor tissues. Sequential gating steps were applied to identify live CD45<sup>+</sup>NKp46<sup>+</sup> NK cells from tumor single-cell suspensions.

**Table S1. Primer sequences used in this study**

| primer | sequence |
| --- | --- |
| fnrS-ompA-*Sp*NA-F | caggccatggagttgttcttattggtggtgttg |
| fnrS-ompA-*Sp*NA-R | gctggcgtttttccataggctccgccccctgacgag |
| lacUV5-HER2-AIDA-F | cctatggaaaaacgccagcaacgcggccttttttacggttcc |
| lacUV5-HER2-R | ttattgttaattaagatctcgaggccgggaagctagagtaag |
| Axe/Txe-F | aattaacaataactagcataaccccttggggcctctaaacg |
| Axe/Txe-R | aagaacaactccatggcctggggtgcctaatgagtgagaatccatgg |
| pACYC-F | atgacaagtaactcgagatcttaattaacaataactag |
| pACYC-R | cggagtttgtttcctttcaccccttatttatttctataaaacaattg |
| hok/sok-F | gggtgaaaggaaacaaactccgggaggcagcgtgatg |
| hok/sok-R | cttccgattggacaacatcagcaaggagaaaggggctac |
| cluster-F | ctgatgttgtccaatcggaagtaaggaaagggtcagaaac |
| cluster-R | aagatctcgagttacttgtcatcgtcgtccttgtaatc |
| EcoRI-*Sp*NA-F | aaggaattcATGCCTGAAGGAGCGGCTTTAAC |
| SaII-*Sp*NA-R | aaagtcgacatctttgctcaaaaagtcccaattaaattttctaaatgata |
| *Sp*NA_E647A-F | aaaatacagcatcaacggtggtacaactaaacaatg |
| *Sp*NA_E647A-R | caccgttgatgctgtattttgcgcacgtctattgttcatc |
| pEgRNA_N20_F | aaacaaagtgcatgttcagggttttagagctagaaatagcaagttaaaataaggc |
| pEgRNA_N20_R | tggatcccctgccagcaaaaggccaggaaccgtaaaaaggccgcgttgc |
| Upstream-F | cttttgctggcaggggatccattttcagccttggcac |
| Upstream-R | atagcggttactttggaagattgtagcgccagtcacagaaaaatgtg |
| Downstream-F | tcttccaaagtaaccgctataaagcgtggcgcggttaatc |
| Downstream-R | aaagaacatgcgcggggaattagtgatggagttttaggac |
| pEgRNA_2_F | attccccgcgcatgttctttcctgcgttatcccctg |
| pEgRNA_2_R | cctgaacatgcactttgtttgctagcattatacctaggactgagctagctgtc |
